# Tandem-Cleavage Linkers Improve the In Vivo Stability and Tolerability of Antibody-Drug Conjugates

**DOI:** 10.1101/2021.01.19.427340

**Authors:** Stepan Chuprakov, Ayodele O. Ogunkoya, Robyn M. Barfield, Maxine Bauzon, Colin Hickle, Yun Cheol Kim, Dominick Yeo, Fangjiu Zhang, David Rabuka, Penelope M. Drake

## Abstract

Although peptide motifs represent the majority of cleavable linkers used in clinical-stage antibody-drug conjugates (ADCs), the sequences are often sensitive to cleavage by extracellular enzymes, such as elastase, leading to systemic release of the cytotoxic payload. This action reduces the therapeutic index by causing off-target toxicities that can be dose-limiting. For example, a common side-effect of ADCs made using peptide-cleavable linkers is myelosuppression, including neutropenia. Only a few reports describe methods for optimizing peptide linkers to maintain efficient and potent tumor payload delivery while enhancing circulating stability. Herein, we address these critical limitations through the development of a tandem-cleavage linker strategy, where two sequential enzymatic cleavage events mediate payload release. We prepared dipeptides that are protected from degradation in the circulation by a sterically-encumbering glucuronide moiety. Upon ADC internalization and lysosomal degradation, the monosaccharide is removed and the exposed dipeptide is degraded, liberating the attached payload inside the target cell. We used CD79b-targeted monomethyl auristatin E (MMAE) conjugates as our model system, and compared the stability, efficacy, and tolerability of ADCs made with tandem-cleavage linkers to ADCs made using standard technology with the vedotin linker. The results—where rat studies showed dramatically improved tolerability in the hematopoietic compartment—highlight the role that linker stability plays in efficacy and tolerability, and offer a means of improving an ADC’s therapeutic index for improved patient outcomes.

## INTRODUCTION

Antibody-drug conjugates (ADCs) are coming into their own as oncology therapeutics. Nine products have been approved for use across a range of cancer indications, including hematological and solid tumors.^1^ The drugs are being used both as monotherapies and—in some cases—as part of combination treatments, highlighting their versatility and utility in the therapeutic armamentarium. Interestingly, the nine approved ADCs vary greatly in terms of their make up, encompassing six different payloads, six distinct linkers, and two conjugation chemistries—lysine-based and cysteine-based. In spite of the differences among the nine approved ADCs, one strong commonality emerges: nearly eighty percent of the drugs use a cleavable linker for tumor payload delivery.

As this data point indicates, cleavable linkers work effectively against a broad variety of target antigens.^2^ For most cleavable linkers, payload release can be triggered early in the endocytic pathway, making lysosomal delivery of the target antigen less of a requirement for ADC activity. This flexibility enables use of a broader range of target antigens, including those that might readily internalize upon ligation but then recycle back to the cell surface rather than progress to the lysosome.^3^ In addition, cleavable linkers enable the use of Payloads that do not tolerate modification and must be in their free form for optimal potency.

One drawback with respect to cleavable linkers is the possibility of payload shedding during ADC circulation prior to tumor targeting. Early, systemic release of payload can reduce the potency of the remaining circulating ADC and can lead to off-target, potentially dose-limiting toxicities. Both complications impact ADC drug development. The former affects interpretation of anti-tumor efficacy in preclinical studies conducted in mice^4^, and the latter limits ADC dosing—and thus potential therapeutic benefit—in patients.^5^ Traditional release mechanisms for cleavable linkers include protonolysis, disulfide reduction, and proteolytic degradation. The latter is most notably embodied by the valine-citrulline (Val-Cit)-PABC linker technology developed by Seagen. The Val-Cit dipeptide motif is designed to be a substrate for cathepsin B and related enzymes in tumor lysosomes. The Val-Cit-PABC linker coupled to a *monomethyl auristatin E* (MMAE) payload (vcMMAE, Figure 1A) is employed on three of the nine approved ADCs and has been tested on more than twenty clinical-stage molecules.^6^

**Figure 1.**
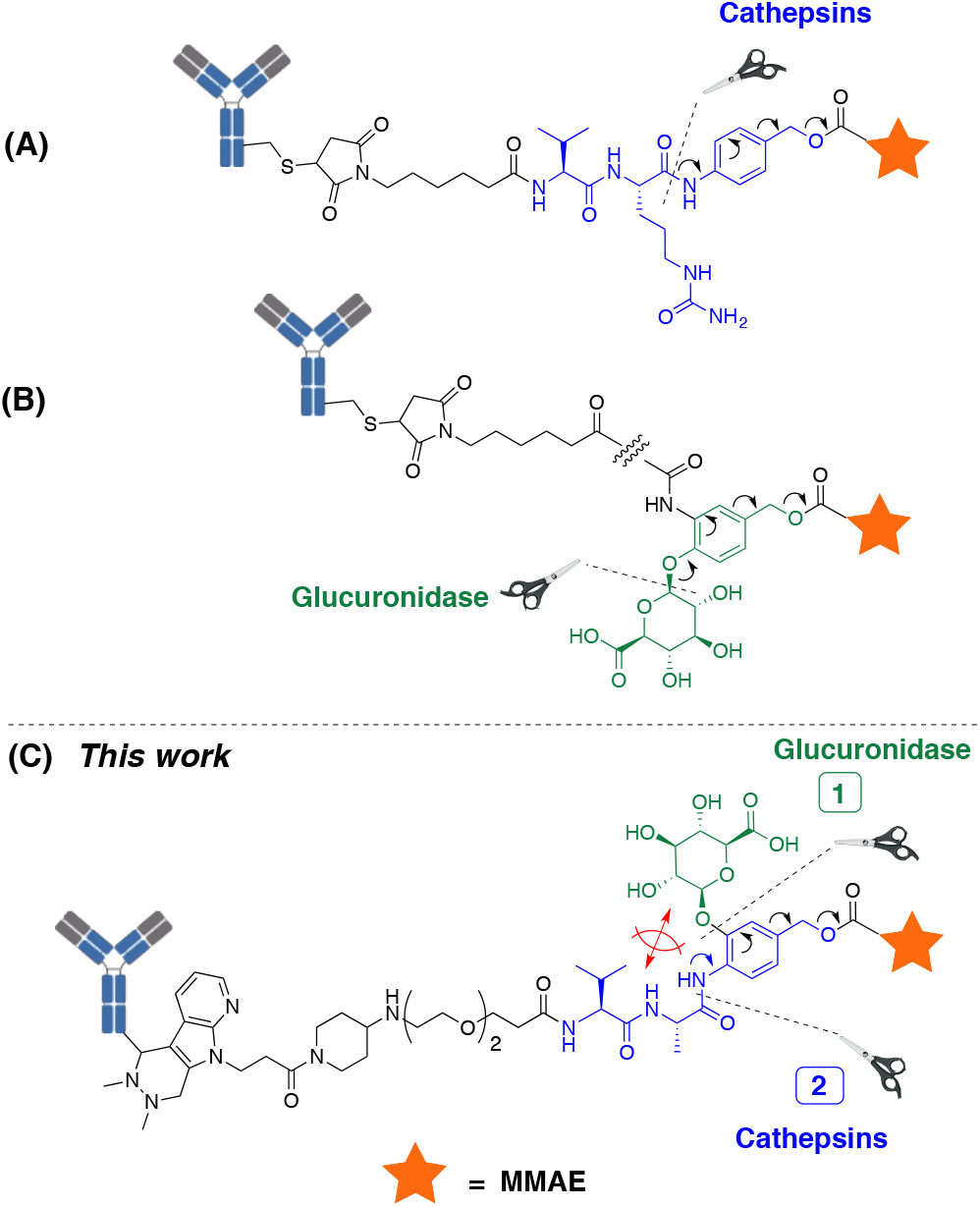
Tandem-cleavage linkers require two sequential enzymatic steps to liberate payload. (A) Cathepsin-cleavable Val-Cit-PABC linker used in vcMMAE conjugates. (B) Glucuronidase-cleavable linker releasing MMAE. (C) Tandem-cleavage linker presented in this study.

A common side-effect of vcMMAE-conjugated ADCs regardless of target antigen is myelosuppression, including neutropenia, thrombocytopenia, and anemia. This off-target toxicity is driven, at least in part, by premature cleavage of the Val-Cit dipeptide via extracellular enzymes encountered during circulation.^7^ In an illuminating study, Zhao and colleagues demonstrate that serine proteases, including serine elastase secreted in part by differentiating neutrophils, can liberate payload from vcMMAE ADCs leading to death of bone mar-row-resident neutrophil precursors.^8^ The data suggest that a cleavable linker resistant to this type of degradation could improve ADC tolerability with respect to bone marrow-derived cells. A number of groups have dedicated research efforts to make cleavable linkers more stable in circulation. Approaches have included using peptide sequences that act as more selective protease substrates ^9–12^, as well as exploiting alternative enzyme classes as release mechanisms.^13,14^

We began with the hypothesis that a linker requiring tandem enzymatic cleavage events, where the second cleavage is hindered until the first cleavage occurs, would limit payload loss during circulation and reduce off-target toxicities. We were inspired by prodrug approaches employing hydrophilic glucuronide moieties,^4,15^ and by glucuronide-containing cleavable linkers (Figure1B),^5,13^ both of which are responsive to β-glucuronidase, a lysosomal enzyme that is often upregulated in malignant cells. We envisioned that a β-glucuronide moiety could serve as a temporary hydrophilic protecting group for dipeptide linkers. The appended monosaccharide would be stable in the circulation but removed upon internalization and lysosomal degradation, thus liberating the dipeptide for further processing (Figure 1C). Herein, we report the development and application of tandem-cleavage linkers for ADCs, and demonstrate that ADCs bearing this novel linker strategy show excellent plasma stability and enhanced tolerability, while remaining efficacious against *in vivo* tumor models.

## RESULTS AND DISCUSSION

### Tandem-cleavage linker design and synthesis

As our goal was to modify a dipeptide linker with a glucuronide protecting group that could be enzymatically-removed inside target cells, we began with an exploration of the chemical space around the dipeptide. To this end, we devised dipeptide linkers modified with glucuronide at positions P1’and P3 (Table 1). Both linkers carried the *MMAE* payload and the Hydrazino-*iso*-Pictet-Spengler (HIPS) conjugation element^6,16^ for ligation to antibodies containing an aldehyde functional group.^7,17^ The P1’ modification was achieved using the Val-Ala dipeptide^8,18^ connected to the PABC self-immolative spacer carrying a pendant *ortho*-glucuronide (construct **1**, Table 1). The P3 modification was achieved by incorporating a glucuronide moiety onto a salicylic acid unit that was attached *N*-terminally to a Val-Cit dipeptide linker (**2**, Table 1).^19,20^ To enable comparison of constructs **1** and **2** with glucuronide-free linkers, control compounds **3** and **4**, respectively, were added to the study (Table 1). Compound **5** is vedotin, the cysteine-reactive vcMMAE construct that has been most commonly used in clinically-tested ADCs and is in the marketed drugs Polivy, Adcetris, and Padcev.^10,21–23^

**Table 1.**
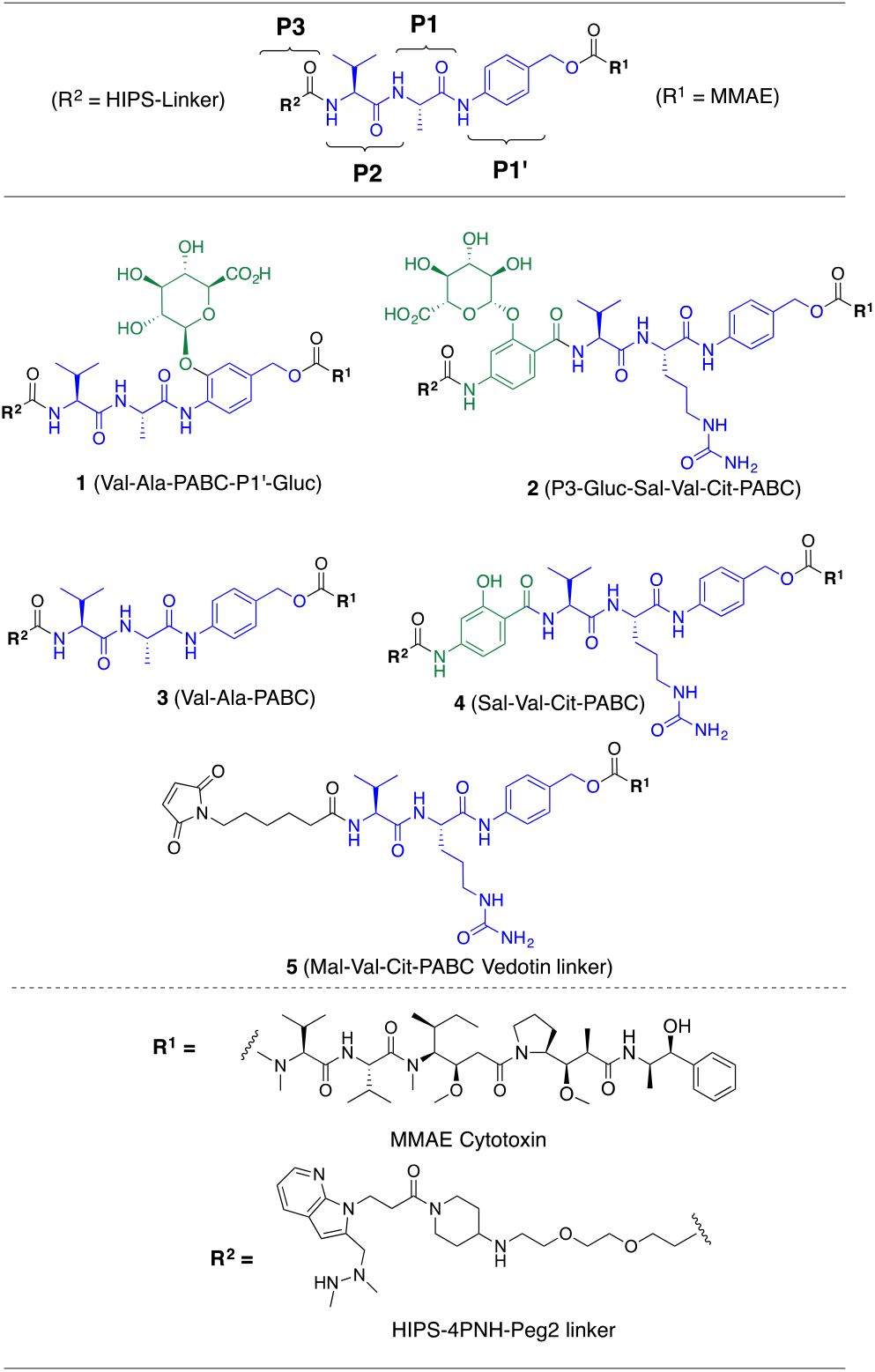
Glucuronide-modified dipeptide linkers and control molecules used in the study

To assemble the P1’-glucuronide portion of compound **1**, the required glycoside bond was formed by reacting acetate-protected glucuronic acid methyl ester bromide **6** with 3-hydroxy-4-nitrobenzaldehyde **7** in the presence of silver(I) oxide (Scheme 1, eq. 1), followed by catalytic hydrogenation of product **8** that affected both the nitro- and carbaldehyde groups and generated intermediate **9**. The latter was further elaborated by coupling the aniline amino group with Boc-protected L-alanine to give **10**. Removal of the Boc group in **10** under acidic conditions, followed by the coupling of the resulting amine with Fmoc-L-valine afforded dipeptide compound **11**, which was further transformed into *p*-nitrophenyl carbonate by engaging the pendant benzyl hydroxy group. Next, activated ester **12** was allowed to react with the free amine of MMAE to furnish the carbamate linkage. After sub-sequent removal of the Fmoc-group, free amine **14** was coupled with pre-assembled Fmoc-protected HIPS-linker **15**. Sub-sequent basic hydrolysis resulted in global deprotection of the glucuronide moiety, removing all four acetate groups and cleaving the methyl ester. Concurrently, the two Fmoc-groups were cleaved under these reaction conditions, liberating the dimethylhydrazine functionality in the HIPS conjugation unit and affording compound **1** in good overall yield (Scheme 1, eq. 1).

**Scheme 1.**
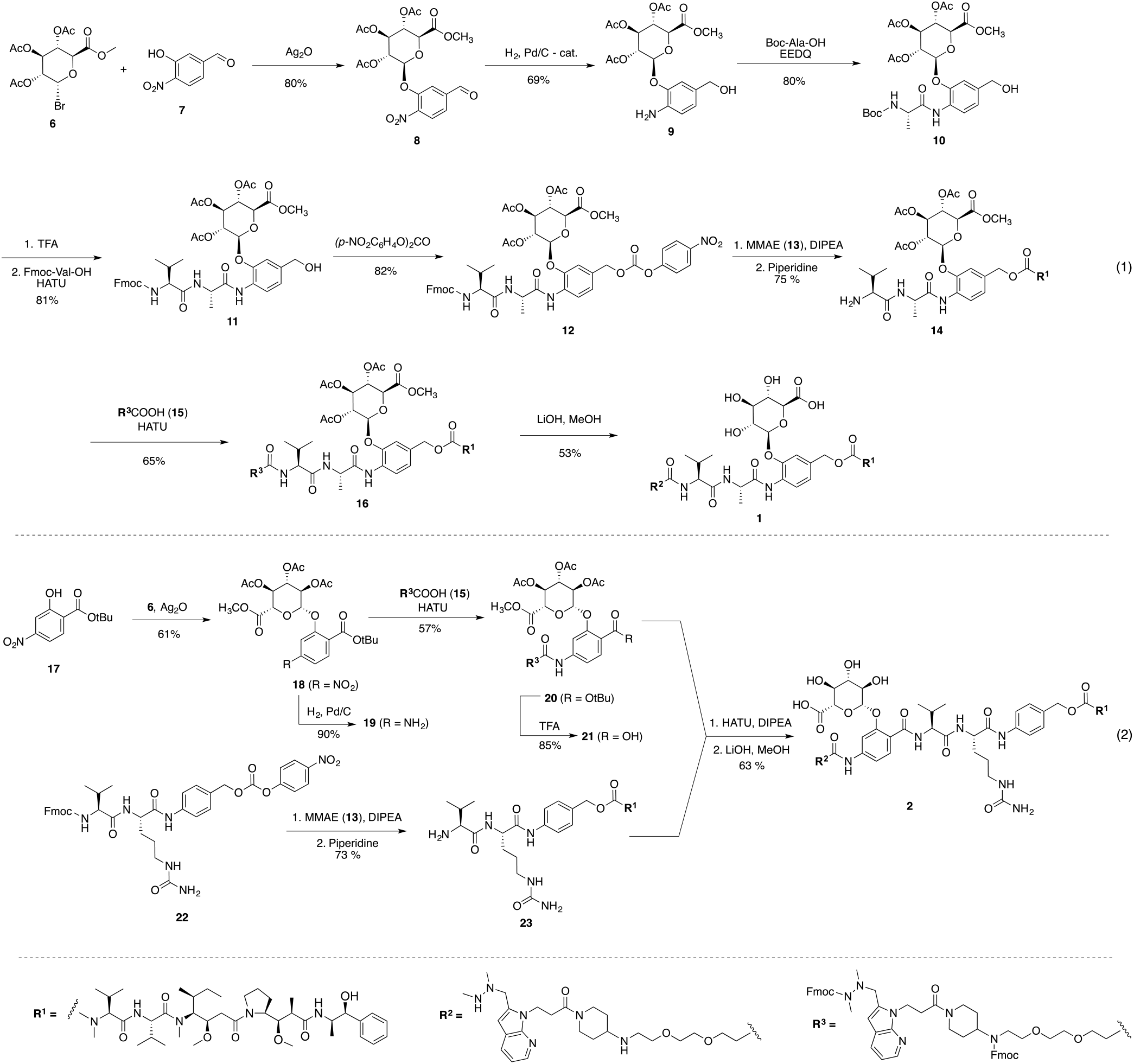
Synthesis of P1’- and P3-glucuronide linker-payloads used in the study

Synthesis of the P3-glucuronide construct **2** originated from the *tert*-butyl ester of *p*-nitrosalicylic acid **17** (Scheme 1, eq. 2). The required glycoside was assembled similarly to the P1’ congener described above with subsequent reduction of the nitro group leading to intermediate **19**. Coupling of the aniline moiety in **19** with HIPS-linker **15**, followed by cleavage of the *tert*-butyl ester, delivered glucuronide assembly **21**. The dipeptide portion of the linker attached to MMAE cytotoxin (**23**, Scheme 1, eq. 2) was prepared from commercially available Fmoc-Val-Cit-PABC-PNP carbonate **22** by forming the carbamate linkage directly followed by the Fmoc group removal. Final synthetic steps included amide coupling between intermediates **21** and **23**, followed by global deprotection under basic hydrolysis conditions to afford the desired construct **2** in good overall yield (Scheme 1, eq. 2). Glucuronide-free constructs **3** and **4** were synthesized in similar fashion starting from the corresponding dipeptides (see Supporting Information for details).

### ADC generation and in vitro testing

Anti-CD79b antibodies^24^ were used as a test system for evaluation of the linker variants shown in Table 1. For constructs **1**-**4** that contained a HIPS conjugation moiety^11,16^, ADCs were generated using antibodies bearing the aldehyde tag at the heavy chain *C*-terminus. Construct **5**, the control maleimide-vcMMAE vedotin linker, was conjugated to wild-type (untagged) anti-CD79b antibody. The drug-to-antibody ratio (DAR) for the HIPS conjugates averaged 1.7 across different constructs, while the vedotin conjugate **5** was generated with an average DAR of 3.2. Exact DAR values are listed in Supplemental Table 1. ADCs were named by listing the linker number (Table 1) followed by the target antigen; for example, the anti-CD79b ADC carrying linker **1** was designated **1**-CD79b.

ADC in vitro potency was tested using the CD79b+ JeKo-1 cell line, derived from a B-cell non-Hodgkin lymphoma (Figure 2A). Cells were treated with serially diluted ADC and cell viability was assessed after 5 days using CellTiter-Glo^®^ to detect ATP. All tested ADCs were equipotent, indicating that the tandem-cleavage requirement did not inhibit efficient Payload release in target cells. ADC serum stability was checked by incubating constructs at 37 °C in rat serum for up to one week (Figure 2B). As anticipated, up to 20% payload loss was observed from conjugates bearing mono-cleavage linkers, such as vcMMAE (**5**) or the non-glycosylated control construct, **3**. Furthermore, tandem-cleavage linkers **1** and **2**, bearing glucuronide at the P1’ and P3 positions, respectively, both yielded stable conjugates with no payload loss detected over 7 days. Surprisingly, the non-glycosylated P3 control linker (**4**) carrying a salicylic acid unit *N*-terminal to the Val-Cit dipeptide was equally stable to the glycosylated versions, showing no payload loss over 7 days. The latter observation was consistent with published reports that modifications *N*-terminal to a dipeptide unit can improve stability.^11^

**Figure 2.**
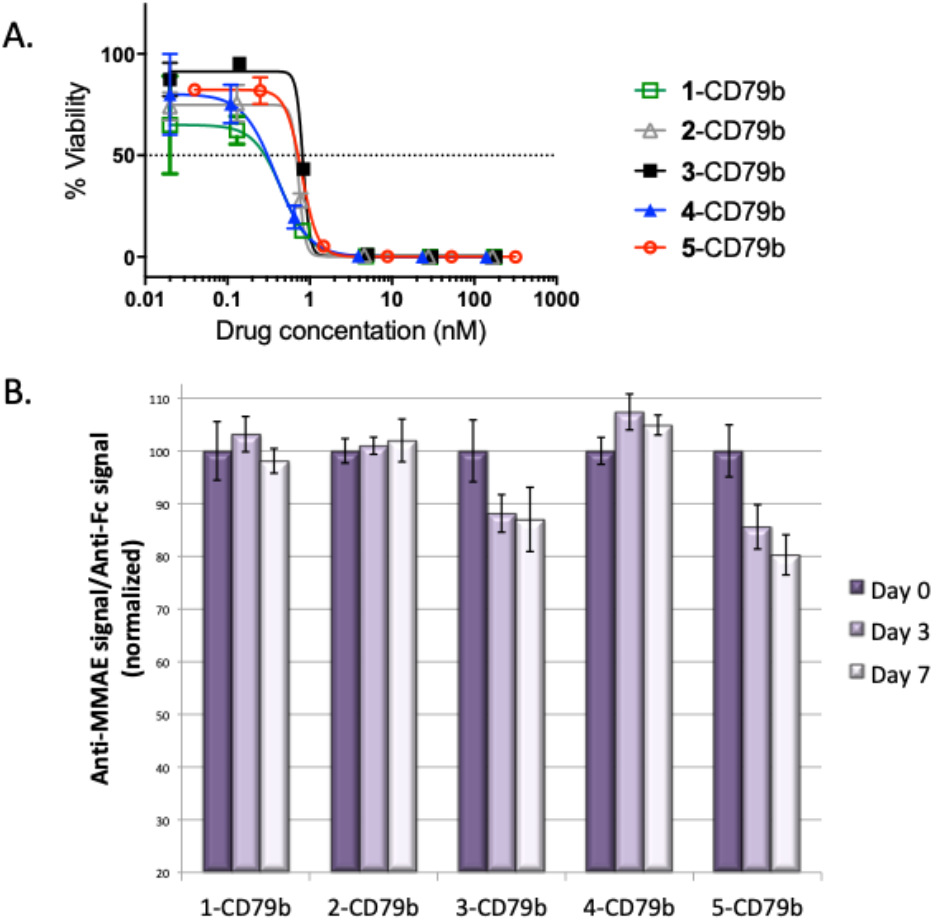
ADCs made with tandem-cleavage linkers are equally potent but generally more stable as compared to ADCs made with mono-cleavage linkers. Anti-CD79b antibodies were conjugated to the linkers shown in Table 1 using either HIPS or maleimide chemistry. (A) The resulting ADCs were tested for in vitro potency against the CD79b+ Jeko-1 cell line using CellTiter Glo to determine cell viability. n = 2; error bars indicate S.D. (B) An ELISA-based assay was used to monitor conjugate stability in rat serum at 37 °C over 7 days. n = 4; error bars indicate S.E.M.

### In vivo efficacy studies

Based on the in vitro results, we elected to take a subset of the conjugates forward into in vivo efficacy studies. First, we used the non-Hodgkin lymphoma xenograft model, Jeko-1, (Figure 3) to compare the efficacy of ADCs bearing tandem-cleavage linkers modified at either the P1’ (1) or P3 positions (2). Mice bearing subcutaneous xenografts received a single intravenous bolus dose of conjugate once tumor volumes reached an average of 167 mm3. While both constructs were efficacious in this model, the ADC bearing linker 1 modified at the P1’ position was superior. This outcome may have reflected the in vitro observation that even the unmodified P3 control construct (5) was stable (Figure 2B), suggesting that payload release might be slower for constructs containing the salicylic acid unit N-terminal to the Val-Cit.

**Figure 3.**
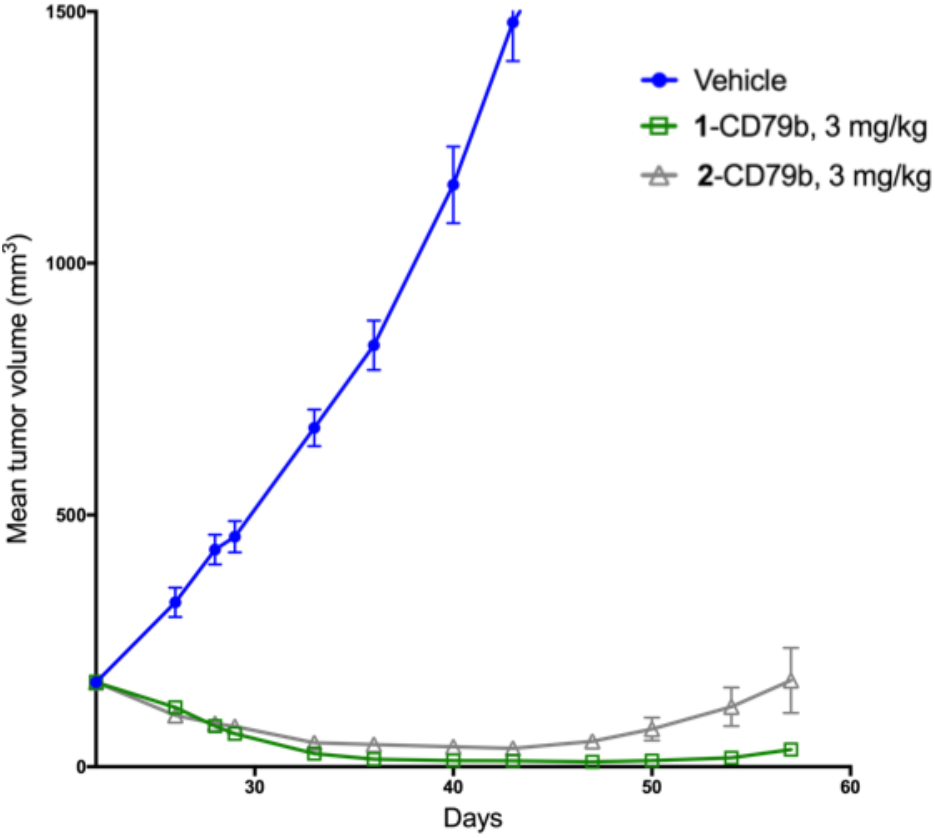
The ADC carrying the P1’ tandem-cleavage linker out-performs the ADC bearing the P3 tandem-cleavage linker against a Jeko-1 xenograft. Mice bearing subcutaneous Jeko-1 tumors were given a single 3 mg/kg intravenous dose of vehicle alone, 1-CD79b, or 2-CD79b. Tumor size was measured twice weekly; animals were monitored for 8 wk post-dose. n = 5 mice/group; error bars indicate S.E.M.

Together, the in vitro and in vivo data pointed to the P1’ position as the optimal site for glucuronide placement in our designs. Next, to measure the impact of the tandem-cleavage approach relative to controls, we used the non-Hodgkin lymphoma Granta-519 xenograft model to test in vivo efficacy. We compared the activity of ADCs made with the P1’ tandem-cleavage linker (1), the control mono-cleavage linker (3) or the conventional maleimide-vcMMAE technology (5). To account for the two-fold difference in DAR between the HIPS and maleimide conjugates, we dosed the mice with an equal ADC payload dose (Figure 4). Specifically, when subcutaneous tumor volumes reached an average of 180 mm3, mice received intravenous doses of HIPS conjugates (1-CD79b and 3-CD79b) at 10 mg/kg or the vcMMAE conjugate (5-CD79b) at 5 mg/kg. In this study, the control ADC bearing the HIPS-conjugated mono-cleavage linker had little affect on tumor growth. By contrast, both the vcMMAE conjugate and the tandem-cleavage HIPS conjugate dramatically reduced tumor volumes. Complete responses (no remaining detectable tumor) were observed in 3 out of 6 animals treated with the vcMMAE (5-CD79b) and 6 out of 6 animals treated with the tandem-cleavage ADC (1-CD79b).

**Figure 4.**
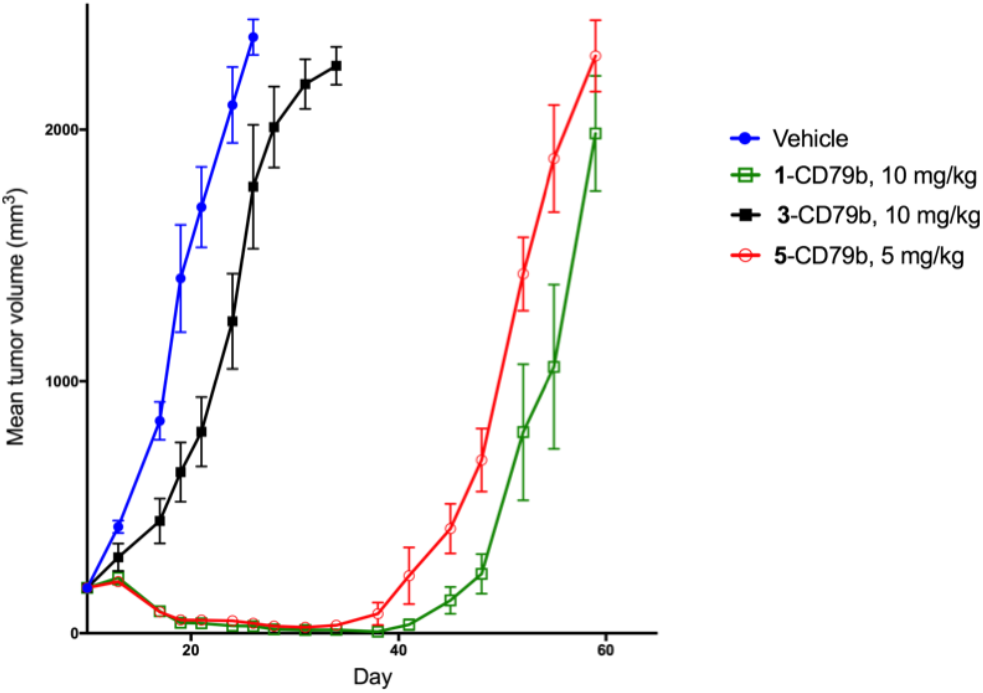
At equal payload doses, the ADC bearing the P1’ tan-dem-cleavage linker provides equal or better efficacy as compared to the vedotin conjugate against a Granta 519 xenograft. Mice carrying Granta 519 xenografts were given a single intravenous dose of vehicle alone, or either 5 or 10 mg/kg of the maleimide or HIPS-conjugated ADC, respectively. Tumor volumes were measured twice weekly, and the study was monitored for 8 wk post-dose. Three of six mice in the 5-CD79b group, and six of six mice in the 1-CD79b group showed complete responses during the study. n = 6 mice/group; error bars indicate S.E.M.

### In vivo tolerability assessment and toxicokinetics

With confirmation of ADC efficacy in-hand, we set out to test the tolerability of the tandem-cleavage ADC as compared to mono-cleavage ADCs in a rat model. We dosed Sprague-Dawley rats once (on Day 1) with an intravenous bolus dose of either 20 mg/kg of the vcMMAE conjugate (5-CD79b) or 40 mg/kg (an equal payload dose) of the tandem-cleavage (1-CD79b) or mono-cleavage (3-CD79b) HIPS conjugates. Animals were monitored for clinical observations and body weight for 11 days post-dose. Clinical chemistry and hematology were assessed on study days 5 and 12. As the anti-CD79b antibody does not recognize rat protein, this study was designed to look at off-target ADC effects. Based on known toxicities of vedotin-conjugates, we anticipated that the major observations would be in the hematopoietic compartment. Indeed, by day 5, animals treated with mono-cleavage conjugates (3-CD79b or 5-CD79b) had marked reductions in circulating neutrophils, monocytes, and eosinophils (Figure 5). By contrast, animals treated with the tandem-cleavage ADC (1-CD79b) showed no evidence of myelosuppression, with white blood cell population counts similar to those of the vehicle control group.

**Figure 5.**
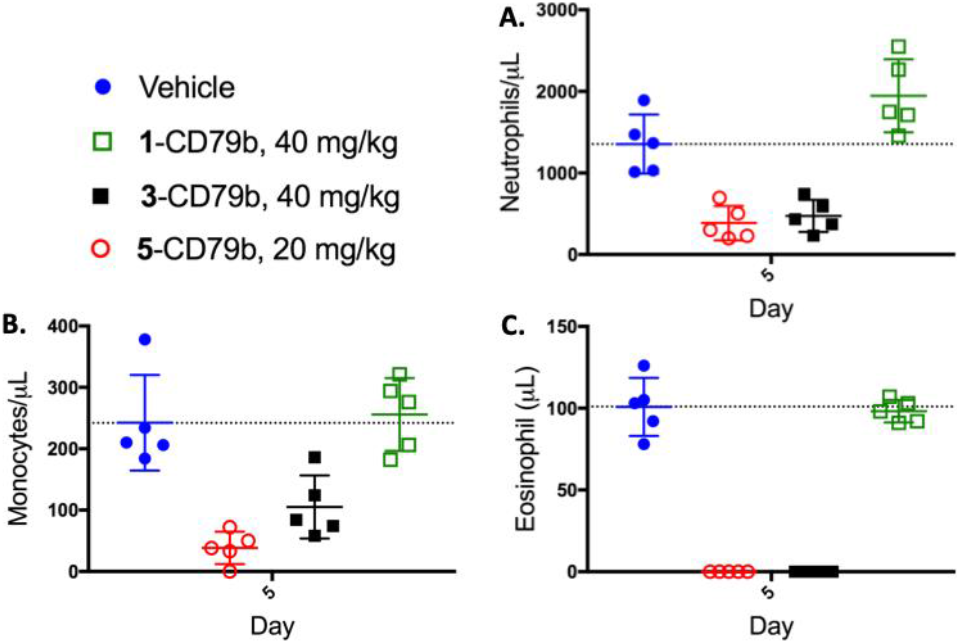
At equal payload doses, the ADC carrying the P1’ tandem-cleavage linker showed improved tolerability relative to ADCs carrying mono-cleavage linkers, with no evidence of myelosuppression. Sprague-Dawley rats (5/group) were given a single intravenous dose of vehicle alone, or ADCs conjugated through HIPS chemistry to a tandem-cleavage (1-CD79b) or mono-cleavage linker (3-CD79b), or through maleimide chemistry to the mono-cleavage vedotin linker (5-CD79b). The HIPS-conjugated ADCs were dosed at 40 mg/kg, while the maleimide conjugate was dosed at 20 mg/kg, resulting in equal payload dosing across the experiment. Circulating (A) neutrophil, (B) monocyte, and (C) eosinophil populations were analyzed at Day 5 post-dose. n = 5; line denotes the mean; error bars indicate S.D.

To confirm dose levels and monitor test article clearance, plasma samples were obtained from the rats at four time points across the tolerability study. ELISA methods were used to measure total antibody and total ADC (conjugated payload) in the samples (Figure 6). The data showed rapid payload loss from both of the mono-cleavage conjugates (3-CD79b and 5-CD79b) while the tandem-cleavage conjugate (1-CD79b) remained mostly intact through day 12. These results provided a rationale for the differences in tolerability among the conjugates. Shed payload from the mono-cleavage conjugates likely drove myelosuppression in animals dosed with those constructs. By contrast, the tandem-cleavage construct was stable in the circulation, retaining payload on the antibody and improving ADC tolerability.

**Figure 6.**
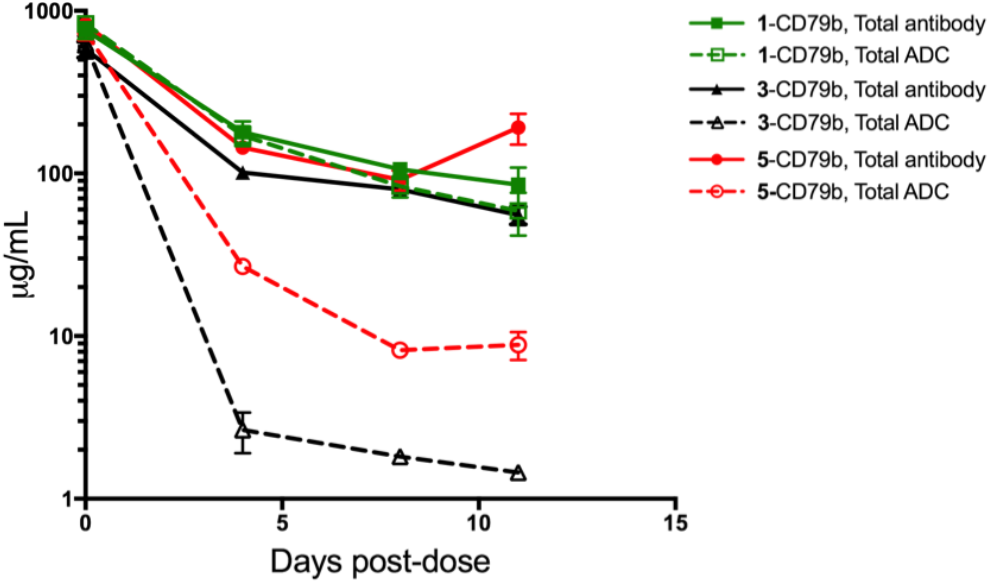
The P1’ tandem-cleavage linker offers greater in vivo stability, with improved payload retention on the antibody, relaive to mono-cleavage linkers such as the vcMMAE vedotin construct. Plasma samples were obtained at four time points post-dose from the rats used in the toxicity study shown in Figure 5. ELISA-based methods were used to quantify total antibody (solid lines) and total ADC (conjugated payload, dashed lines). n = 5; error bars indicate S.D.

Interestingly, the difference in the rate of payload loss from the two mono-cleavage conjugates (both featuring dipeptide PABC motifs) likely reflects the effect of conjugation site on ADC stability. With the HIPS conjugate placed at the antibody heavy chain C-terminus, the linker was likely exposed as a substrate for extracellular enzymes such as elastase. By contrast, the vcMMAE construct carried linkers at the cysteine residues in the antibody hinge region. These were likely partially shielded by the antibody and somewhat protected from degradative enzymes. Considering the exposed nature of the HIPS conjugation site, the stability obtained using the tandem-cleavage linker is even more remarkable. Collectively, this work highlights both the complexity of ADC structure/function relationships, as well as the opportunities for improving on existing technologies through rational biology-guided design.

In conclusion, we developed a novel tandem-cleavage linker system for use in ADCs. We used the MMAE payload as a model because the abundant preclinical and clinical data generated with the classical vedotin linker conjugates could be used as a benchmark for comparison. Our tandem-cleavage approach yielded conjugates that were stable in the circulation, efficacious against xenograft tumor models, and well-tolerated in rat toxicity studies, with negligible effects on bone marrow-derived cells. Moreover, the tandem-cleavage strategy is transferable to a wide-range of other payloads, and thus offers a generalizable method for developing stable, well-tolerated antibody-drug conjugates.

## EXPERIMENTAL

### General

All animal studies were conducted in accordance with Institutional Animal Care and Use Committee guidelines and were performed at Crown Bioscience or Bolder Bio-PATH. The murine anti-MMAE antibody (B12A2) was purchased from Levena Biopharma. Horseradish peroxidase (HRP)-conjugated secondary antibodies were obtained from Jackson Immunoresearch. The Jeko-1 cell line was purchased from the ATCC cell bank, where it was authenticated by morphology, karyotyping, and PCR-based approaches. The cell line has not been retested since beginning culture in our laboratory five years ago. All original synthetic procedures can be found in the Supporting Information.

### Cloning, expression, and purification of tagged antibodies

Antibodies were generated using standard cloning and purification techniques as previously described.^12,25^ ExpiCHO cells stably overexpressing human formylglycine-generating enzyme were used for transient production of antibodies in media supplemented with 100 μM copper(II) sulfate.^26^

### Bioconjugation and HPLC analytics

#### HIPS-conjugation of aldehyde-tagged antibodies

Antibodies bearing an aldehyde tag at the heavy chain C-terminus (15 mg/mL) were conjugated with 8 mol. equivalents of linker-payload (drug:antibody) for 72 h at 37° C in 50 mM sodium citrate, 50 mM NaCl pH 5.5 containing 0.85-4.25% DMA. Free drug was removed using tangential flow filtration. To determine the DAR of the final product, ADCs were examined by analytical HIC (Tosoh #14947) with mobile phase A: 1.5 M ammonium sulfate, 25 mM sodium phosphate pH 7.0, and mobile phase B: 25% isopropanol, 18.75 mM sodium phosphate pH 7.0. To determine aggregation, samples were analyzed using analytical size exclusion chromatography (SEC; Tosoh #08541) with a mobile phase of 5% isopropanol, 300 mM NaCl, 25 mM sodium phosphate pH 6.8.

#### Maleimide conjugation of untagged (wild-type) antibodies

Antibodies (5 mg/mL) were reduced using 2.5 mol. equivalents of TCEP for 90 min at 37° C in in PBS, pH 8.0, 1 mM DTPA. TCEP was removed and the protein was exchanged into PBS, pH 7.4, 1 mM DTPA using tangential flow filtration. Reduced antibody (3 mg/mL) was conjugated with 10 mol. equiv of maleimide-Val-Cit-MMAE for 60 min on ice. Free drug was removed and final ADC was exchanged into PBS, pH 7.4 using tangential flow filtration.

#### In vitro cytotoxicity assays

Cell lines were plated in 96-well plates (Costar 3610) at a density of 5 x 104 cells/well in 100 μL of growth media. The next day cells were treated with 20 μL of test articles serially-diluted in media. After incubation at 37 °C with 5% CO2 for 5 days, viability was measured using the Promega CellTiter Glo® reagent according to the manufacturer’s recommendations. GI50 curves were calculated in GraphPad Prism normalized to the payload concentration.

#### In vitro stability

ADCs were spiked into rat serum at 50 μg/mL. Samples were aliquoted and stored at −80 °C until use. Aliquots were placed at 37°C under 5% CO2 for the indicated times, and then were analyzed by ELISA. A freshly thawed aliquot was used as a reference starting value for conjugation. Analytes were diluted 1:1000 into casein blocking buffer (Thermo Scientific), so that the concentrations would be in the linear range of the ELISA assays. Samples were measured together on one plate to enable comparisons across time points. Analytes were captured on plates coated with an anti-human Fab-specific antibody (Jackson Immunoresearch). Then, conjugated payload was detected with an anti-MMAE antibody (Clone B12A2, Levena Biopharma) followed by an HRP-conjugated anti-mouse secondary antibody. Total antibody was detected with an HRP-conjugated anti-human Fc-specific secondary antibody (Jackson Immunoresearch). Bound secondary antibody was visualized using Ultra TMB One-Step ELISA substrate (Thermo Fisher). The colorimetric reaction was stopped with H2SO4, and absorbance at 450 nm was determined using a Molecular Devices SpectraMax M5 plate reader. Data analysis was performed in Excel. Each sample was analyzed in quadruplicate, and the average values were used. The ratio of anti-MMAE signal to anti-Fc signal was used as a measure of antibody conjugation.

#### Xenograft studies

Female NOD/SCID or CB17/SCID mice were inoculated subcutaneously with 10e6 Granta 519 or JeKo-1 cells, respectively. The JeKo-1 cells were implanted in a suspension of Matrigel. Tumors were measured twice weekly and tumor volume was estimated according to the formula: tumor volume (mm3)=w2xl2 where w = tumor width and l = tumor length. When tumors reached the desired mean volume, animals were randomized into groups of 5-6 mice and were dosed as described in the text. Animals were euthanized at the end of the study or when tumors reached 2000 mm3.

#### Rat toxicology study

Male Sprague-Dawley rats (5/group) were given a single intravenous dose of vehicle alone, 20 mg/kg of 6-CD79b ADC (the maleimide conjugate), or 40 mg/kg of 1-CD79b or 4-CD79b ADCs (the aldehyde tag conjugates). Animals were observed for 12 days post-dose. Body weights were recorded on days 0, 1, 4, 8, and 11. Blood was collected from all animals at 8 h and at 5, 9, and 12 d and was used for toxicokinetic analyses (all time points) and for clinical chemistry and hematology analyses (days 5 and 12).

#### Toxicokinetic sample analysis

The concentrations of total antibody and total ADC in rat toxicokinetic samples from the single dose study were quantified by ELISA as previously described.^27^ For total antibody, conjugates were captured with an anti-human IgG-specific antibody and detected with an HRP-conjugated anti-human Fc-specific antibody. For total ADC, conjugates were captured with an anti-human Fab specific antibody and detected with a mouse anti-MMAE primary antibody (Levena Biopharma), followed by an HRP-conjugated anti-mouse secondary antibody. Bound secondary antibody was detected using Ultra TMB One-Step ELISA substrate (Thermo Fisher). After quenching the reaction with sulfuric acid, signals were read by taking the absorbance at 450 nm on a Molecular Devices Spectra Max M5 plate reader equipped with SoftMax Pro software. Data were analyzed using GraphPad Prism and Microsoft Excel software.

## ASSOCIATED CONTENT

### Supporting Information

Full experimental procedures, characterization data, and NMR spectra. The Supporting Information is available free of charge on the ACS Publications website.

## AUTHOR INFORMATION

### Author Contributions

The manuscript was written through contributions of all authors. / All authors have given approval to the final version of the manu-script.

## Notes

### Competing Interest Statement

All authors are employees of Catalent Biologics.

