## Supplemental Info for "Tandem-Cleavage Linkers Improve the In Vivo Stability and Tolerability of Antibody-Drug Conjugates"

Catalent Pharma Solutions, 5959 Horton Street, Suite 400, Emeryville, California 94608, United States

#### Contents

General Information…………………………………………………………… SI 2

Experimental Procedures……………………………………………………………. SI 3

Synthesis of P1'-Glucuronide MMAE construct **1**……………………………………... SI 3

Synthesis of P3-Glucuronide MMAE construct **2**………………………………………. SI 7

Synthesis of control MMAE construct **3**…………………………………………………. SI 9

Synthesis of control MMAE construct **4**…………………………………………………. SI 10

Synthesis of AzaHIPS-4PNH-PEG2 linker **15**…………………………………………… SI 12

NMR spectra…………………………………………………………………………….... SI 14

Table SI-1. DARs of ADCs used in the reported studies………………………………… SI 22

#### General Information

Synthetic reagents were purchased from Sigma-Aldrich, Acros Organics, AK Scientific, or other commercial sources and used without purification. Anhydrous solvents were obtained from Sigma-Aldrich in sealed bottles and stored under nitrogen. Monomethyl auristatin A (MMAE, **13**) was purchased from DC Chemicals or BroadPharm and used as received. Synthesis of Fmoc-AzaHIPS-COOH **31** was previously described.[1] Maleimide-MMAE compound **5** was obtained from commercial sources or synthesized using literature procedures.[2] Compounds **22**, **24**, and **31** were obtained commercially from Shanghai Medicilon and used without purification. Flash chromatography was performed using a Biotage Isolera chromatography system. Preparative HPLC purifications were performed using a Waters preparative HPLC unit equipped with a Phenomenex Kinetex 5 μm EVO C18 150 x 21.2 mm column. Low-resolution mass spectra (LRMS) were acquired on an Agilent Technology 6120 Quadrupole LC/MS, equipped with an Agilent 1260 Infinity HPLC system, G1314 variable wavelength detector, and Agilent Poroshell 120 SB C18, 4.6 mm x 50 mm column at room temperature using a 10-100% gradient of water and acetonitrile containing 0.1% formic acid. High-resolution mass spectra (HRMS) were acquired on a Thermo Scientific Q Exactive mass spectrometer (70,000 resolution, positive polarity 300 to 2100 m/z scan range), equipped with an Agilent 1260 Infinity HPLC system and Zorbax RRHD Eclipse Plus C18 2.1 x 100 mm 1.8 μm column. $^{1}$H and $^{13}$C NMR spectra were recorded on a 600 MHz Bruker Avance III spectrometer using a Prodigy Cryoprobe. Chemical shifts (δ) are reported as parts per million (ppm), residual solvent peaks are used as references to convert chemical shifts to the TMS scale.

[1] Albers, A. E.; Garofalo, A. W.; Drake, P. M.; Kudirka, R.; de Hart, G. W.; Barfield, R. M.; Baker, J.; Banas, S.; Rabuka, D. *Eur. J. Med. Chem. 2014*, **88**, 3–9.
[2] Doronina, S. O.; Senter, P. D.; Toki, B. E.; Ebens, A. J.; Kline, T. B.; Polakis, P.; Sliwkowski, M. X.; Spencer, S. D. U.S. Pat. Appl. Publ. (2005), US 20050238649 A1.

#### Experimental Procedures

##### Synthesis of P1'-Glucuronide MMAE construct 1

*(2S,3R,4S,5S,6S)-2-(5-formyl-2-nitrophenoxy)-6-(methoxycarbonyl)tetrahydro-2H-pyran-3,4,5-triyl triacetate (**8**).* To an oven dried 50 mL oven-dried round bottom flask were added 3-hydroxy-4-nitrobenzaldehyde (**7**, 334 mg, 2.0 mmol), 2.38 g (6.0 mmol) of acetobromo-α-D-glucuronic acid methyl ester (**6**), and 20 mL of anhydrous acetonitrile. The resulting mixture was treated with silver (I) oxide (3.7 g, 16 mmol) and stirred vigorously at room temperature in the dark for 24 hours. Reaction mixture was diluted with 20 mL of ethyl acetate, filtered through a pad of silica gel and concentrated under vacuum. The residue was purified on silica gel, eluted with 0-25% ethyl acetate-hexanes gradient to obtain 767 mg (1.6 mmol, 80 % yield) of product **8** as a white foamy solid. $^{1}$H NMR (600 MHz, $CDCl_3$) δ 10.03 (s, 1H), 7.89 (d, *J* = 8.2 Hz, 1H), 7.86 (d, *J* = 1.5 Hz, 1H), 7.71 (dd, *J* = 8.2, 1.5 Hz, 1H), 5.42 – 5.34 (m, 2H), 5.32 (t, *J* = 8.3 Hz, 1H), 5.27 (dd, *J* = 8.2, 6.4 Hz, 1H), 4.31 (d, *J* = 8.8 Hz, 1H), 3.72 (s, 3H), 2.11 (s, 3H), 2.06 (s, 3H), 2.05 (s, 3H). $^{13}$C NMR (151 MHz, $CDCl_3$) δ 189.82, 170.03, 169.40, 169.25, 166.75, 149.49, 144.67, 139.67, 125.71, 125.05, 119.26, 99.24, 72.60, 70.70, 70.07, 68.42, 53.17, 20.67, 20.63, 20.61. LRMS (ESI): *m/z* 506.1 $[M+Na]^+$, Calcd for $C_{20}H_{21}NO_{13}$ *m/z* 506.1.

*(2S,3R,4S,5S,6S)-2-(2-amino-5-(hydroxymethyl)phenoxy)-6-(methoxycarbonyl)tetrahydro-2H-pyran-3,4,5-triyl triacetate (**9**).* In a 25 mL round-bottom flask were mixed compound **8** (670 mg, 1.47 mmol), 40 μL (0.28 mmol) of triethylamine, and palladium on carbon (10 wt. %, 160 mg, 0.14 mmol) in 10 mL of anhydrous ethyl acetate. The mixture was flushed with argon, reaction flask was sealed with a rubber septum, equipped with a hydrogen balloon, and stirred at room temperature for 24 hours. Reaction mixture was filtered through a pad of celite, concentrated under vacuum, and purified on silica gel eluted with 0-75% ethyl acetate-hexane to obtain 458 mg (1.01 mmol, 69% yield) of product as an off-white foamy solid. $^{1}$H NMR (600 MHz, $CDCl_3$) δ 6.94 (d, *J* = 1.8 Hz, 1H), 6.86 (dd, *J* = 8.0, 1.8 Hz, 1H), 6.66 (d, *J* = 8.0 Hz, 1H), 5.37 – 5.24 (m, 3H), 5.02 (d, *J* = 7.5 Hz, 1H), 4.52 – 4.44 (m, 2H), 4.19 (d, *J* = 9.6 Hz, 1H), 3.73 (s, 3H), 3.21 (br s, 2H), 2.05 (s, 3H), 2.04 (s, 3H), 2.04 (s, 3H). $^{13}$C NMR (151 MHz, $CDCl_3$) δ 170.10, 169.83, 169.59, 167.03, 144.30, 137.29, 131.28, 123.59, 115.80, 115.59, 100.36, 72.56, 71.72, 71.10,

69.36, 65.12, 53.10, 20.85, 20.72, 20.59. LRMS (ESI): *m/z* 456.1 $[M+H]^+$, Calcd for $C_{20}H_{25}NO_{11}$ *m/z* 456.1.

*(2S,3R,4S,5S,6S)-2-(2-((S)-2-((tert-Butoxycarbonyl)amino)propanamido)-5-(hydroxymethyl)phenoxy)-6-(methoxycarbonyl)tetrahydro-2H-pyran-3,4,5-triyl triacetate (**10**).* To a mixture of aniline compound **9** (567 mg, 1.25 mmol) and Boc-L-Ala-OH (235 mg, 1.25 mmol) in 5 mL of anhydrous DCM were added EEDQ (370 mg, 1.5 mmol). The resulting mixture was stirred overnight at room temperature in the dark, then purified directly on silica gel, eluted with 0-80% ethyl acetate-hexane gradient to obtain 625 mg (1.0 mmol, 80% yield) of title compound **10** as a white solid. $^1$H NMR (600 MHz, $CDCl_3$) δ 8.26 – 8.21 (m, 2H), 7.00 – 6.96 (m, 2H), 5.46 (s, 1H), 5.42 (t, *J* = 9.5 Hz, 1H), 5.33 (dd, *J* = 9.7, 7.6 Hz, 1H), 5.28 (t, *J* = 9.6 Hz, 1H), 5.17 (d, *J* = 7.6 Hz, 1H), 4.58 (s, 2H), 4.42 (s, 1H), 4.28 (d, *J* = 9.7 Hz, 1H), 3.73 (s, 3H), 2.06 (s, 3H), 2.05 (s, 3H), 2.04 (s, 3H), 1.45 (s, 9H), 1.43 (d, *J* = 7.2 Hz, 3H). $^{13}$C NMR (151 MHz, $CDCl_3$) δ 171.68, 170.69, 169.92, 169.55, 166.79, 155.35, 145.53, 137.42, 127.59, 122.29, 120.86, 112.39, 99.26, 79.98, 72.59, 71.37, 71.21, 69.35, 64.70, 53.23, 51.13, 28.47 (3C), 20.89, 20.67, 20.57, 19.08. LRMS (ESI): *m/z* 627.2 $[M+H]^+$, Calcd for $C_{28}H_{38}N_2O_{14}$ *m/z* 627.2.

*(2S,3R,4S,5S,6S)-2-(2-((S)-2-((S)-2-((((9H-Fluoren-9-yl)methoxy)carbonyl)amino)-3-methylbutanamido)propanamido)-5-(hydroxymethyl)phenoxy)-6-(methoxycarbonyl)tetrahydro-2H-pyran-3,4,5-triyl triacetate (**11**).* Compound **10** (545 mg, 0.87 mmol) was dissolved in a mixture of DCM-TFA (1:1, 4 mL) at ambient temperature, and let stand for 20 minutes. Solvents were removed under vacuum; the yellow oil residue was dried under high vacuum for few hours and subjected to the next step without any purification. LRMS (ESI): *m/z* 527.2 $[M+H]^+$, Calcd for $C_{23}H_{30}N_2O_{12}$ *m/z* 527.2.

Crude amine (~0.87 mmol) was dissolved in 2 mL of anhydrous DMF. To this mixture were added DIPEA (0.46 mL, 2.6 mmol) and HOAt (118 mg, 0.87 mmol). In a separate vial, HATU (330 mg, 0.87 mmol) was mixed Fmoc-L-Val-OH (295 mg, 0.87 mmol) in 2 mL of DMF, stirred at room temperature for 30 minutes and added to the mixture of amine and DIPEA. The resulting mixture was stirred at room temperature for 2 hours, quenched by pouring into 20 mL of saturated aqueous ammonium chloride solution, extracted with ethyl acetate (2x20 mL). After removal of solvents under vacuum the residue was purified on silica gel with 20-80% EtOAc-DCM as eluent to obtain

597 mg (0.71 mmol, 81% yield) of the title compound **11** as a white solid. $^{1}$H NMR (600 MHz, DMSO) δ 8.71 (s, 1H), 8.14 (d, *J* = 6.7 Hz, 1H), 7.94 (d, *J* = 8.2 Hz, 1H), 7.89 (d, *J* = 7.5 Hz, 2H), 7.75 (t, *J* = 7.4 Hz, 2H), 7.44 – 7.37 (m, 3H), 7.32 (tdd, *J* = 7.4, 2.8, 1.1 Hz, 2H), 7.08 (d, *J* = 1.8 Hz, 1H), 7.06 – 6.93 (m, 1H), 5.64 (d, *J* = 7.7 Hz, 1H), 5.52 (t, *J* = 9.6 Hz, 1H), 5.25 (dd, *J* = 9.7, 7.7 Hz, 1H), 5.22 (t, *J* = 5.7 Hz, 1H), 5.10 (t, *J* = 9.7 Hz, 1H), 4.70 (d, *J* = 9.9 Hz, 1H), 4.48 (t, *J* = 6.9 Hz, 1H), 4.45 (d, *J* = 5.6 Hz, 2H), 4.35 – 4.29 (m, 1H), 4.26 - 4.20 (m, 2H), 3.97 (dd, *J* = 9.2, 6.7 Hz, 1H), 3.63 (s, 3H), 2.04 (s, 3H), 2.01 (s, 3H), 1.99 (s, 3H), 1.34 (d, *J* = 7.0 Hz, 3H), 0.89 (d, *J* = 6.7 Hz, 3H), 0.85 (d, *J* = 6.8 Hz, 3H). $^{13}$C NMR (151 MHz, DMSO) δ 171.14, 170.82, 169.85, 169.49, 169.32, 167.03, 156.13, 146.15, 143.86, 143.81, 140.71, 140.68, 139.15, 127.63, 127.61, 127.05, 127.04, 126.94, 125.38, 125.36, 121.29, 121.16, 120.08, 120.07, 113.63, 98.32, 70.98, 70.89, 70.87, 68.92, 65.70, 62.42, 59.78, 52.57, 49.18, 46.68, 30.42, 20.41, 20.29, 20.22, 19.22, 18.00, 17.72. LRMS (ESI): *m/z* 870.3 $[M+H]^+$, Calcd for $C_{43}H_{49}N_3O_{15}$ *m/z* 870.3.

*(2S,3R,4S,5S,6S)-2-(2-((S)-2-((S)-2-((((9H-fluoren-9-yl)methoxy)carbonyl)amino)-3-methylbutanamido)propanamido)-5-((((4-nitrophenoxy)carbonyl)oxy)methyl)phenoxy)-6-(methoxycarbonyl)tetrahydro-2H-pyran-3,4,5-triyl triacetate (**12**).* To a solution of benzyl alcohol **11** (334 mg, 0.39 mmol) and bis(4-nitrophenyl) carbonate (240 mg, 0.79 mmol) in THF (3 mL) was added DIPEA (0.1 mL, 0.57 mmol). Reaction mixture was stirred for 24 hours at room temperature, then solvent was removed under vacuum, and the residue was purified by column chromatography on silica gel (ethyl acetate-hexane, 1:9 to 9:1 v/v) to yield PNP carbonate product **12** (328 mg, 0.32 mmol, 82% yield) as an off-white solid. $^{1}$H NMR (600 MHz, $CDCl_3$) δ 8.39 (d, *J* = 8.3 Hz, 1H), 8.32 (s, 1H), 8.29 – 8.26 (m, 2H), 7.76 (d, *J* = 7.6 Hz, 2H), 7.61 (t, *J* = 6.4 Hz, 2H), 7.42 – 7.34 (m, 4H), 7.31 (t, *J* = 7.5 Hz, 2H), 7.16 (d, *J* = 8.3 Hz, 1H), 7.03 (s, 1H), 6.76 (d, *J* = 7.5 Hz, 1H), 5.53 (d, *J* = 9.0 Hz, 1H), 5.44 (t, *J* = 9.4 Hz, 1H), 5.41 – 5.30 (m, 2H), 5.21 (s, 2H), 5.16 (d, *J* = 7.4 Hz, 1H), 4.80 – 4.72 (m, 1H), 4.45 (dd, *J* = 10.6, 7.3 Hz, 1H), 4.36 (dd, *J* = 10.6, 7.2 Hz, 1H), 4.26 (d, *J* = 9.7 Hz, 1H), 4.23 (t, *J* = 7.2 Hz, 1H), 4.16 (dd, *J* = 9.0, 5.9 Hz, 1H), 3.72 (s, 3H), 2.24 – 2.17 (m, 1H), 2.12 (s, 3H), 2.06 (s, 6H), 1.51 (d, *J* = 7.1 Hz, 3H), 1.02 (d, *J* = 6.7 Hz, 3H), 0.95 (d, *J* = 6.7 Hz, 3H). $^{13}$C NMR (151 MHz, $CDCl_3$) δ 171.25, 171.09, 170.83, 169.95, 169.47, 166.67, 156.58, 155.57, 152.50, 145.59, 145.21, 144.04, 143.96, 141.43, 130.15, 129.23, 127.82, 127.83, 127.20, 125.45, 125.30, 125.25, 124.80, 121.89, 120.89, 120.11, 120.08, 114.57, 99.48, 72.76, 71.44, 70.97, 70.56, 69.28, 67.21, 60.21, 53.26, 49.99, 47.34, 31.68, 21.16,

20.69, 20.59, 19.51, 18.65, 17.70. LRMS (ESI): *m/z* 1013.3 $[M+H]^+$, Calcd for $C_{50}H_{53}N_4O_{19}$ *m/z* 1013.3.

*H-Val-Ala-PABC(o-Gluc-OAc)-MMAE (**14**).* In an oven-dried 20 mL glass scintillation vial were mixed MMAE as a TFA salt (**13**, 150 mg, 0.18 mmol) and PNP carbonate 12 (160 mg, 0.16 mmol) in 2 mL of anhydrous DMF. This mixture was treated with 84 uL (0.48 mmol) of DIPEA and allowed to react at room temperature for 2 hours. DIPEA was removed under vacuum, the residual solution was treated with 32 uL (0.32 mmol) of piperidine at 0 °C for 7 hours, and purified by reversed-phase HPLC (C18, acetonitrile-water 5-95% gradient with 0.05% TFA). Pure fractions were lyophilized to give 160 mg (0.12 mmol, 75% yield over 2 steps) of the title compound **14** as a white powder. LRMS (ESI): *m/z* 1369.8 $[M+H]^+$, Calcd for $C_{68}H_{104}N_8O_{21}$ *m/z* 1369.7.

*Fmoc-AzaHIPS-4PN(Fmoc)-PEG2-Val-Ala-PABC(o-Gluc-OAc)-MMAE (**16**).* To a solution of compound **14** (32 mg, 23 µmol) and HIPS linker **15** (27 mg, 29 µmol) in DMF (1 mL) were added DIPEA (10 µL, 57 µmol) and HATU (11 mg, 29 µmol). The reaction mixture was stirred for 1 hour at room temperature. The mixture was diluted with DCM (70 mL), washed sequentially with saturated aqueous $NH_4Cl$ (50 mL), water (50 mL), and saturated aqueous NaCl (10 mL). Organic layer was dried over $MgSO_4$, filtered, and evaporated. The residue was purified on silica gel (EtOAc-hexane, 10-90% v/v) to yield product **16** (33 mg, 15 µmol, 65% yield) as a white solid. LRMS (ESI): *m/z* 1151.0 $[M+2H]^{++}$, Calcd for $C_{123}H_{164}N_{14}O_{29}$ *m/z* 1150.6.

*AzaHIPS-4PNH-PEG2-Val-Ala-PABC(o-Gluc)-MMAE (**1**).* To a solution of MMAE compound **12** (33 mg, 15 µmol) in THF (1 mL) at 0 °C were added a solution of lithium hydroxide monohydrate (3 mg, 71 µmol) in water (0.25 mL). Reaction mixture was warmed up to room temperature and stirred for 1 hour, then quenched by addition of acetic acid (3 µL, 53 µmol). After solvents were removed under vacuum, the residue was purified by reversed-phase chromatography (C18 column, 5-80% acetonitrile/water/0.05% TFA). Fractions containing the desired compound were pooled and lyophilized to yield P1'-glucuronide-MMAE construct **1** (13 mg, 8 µmol, 53% yiled) as a white solid. HRMS (ESI): *m/z* 1715.9798 $[M+H]^+$, Calcd for $C_{86}H_{135}N_{14}O_{22}$ *m/z* 1715.9870.

**Synthesis of P3-glucuronide-MMAE construct 2**

*tert-Butyl 2-hydroxy-4-nitrobenzoate (**17**).* To a stirred solution of 2-hydroxy-4-nitrobenzoic acid (366 mg, 2.0 mmol) in 5 mL of anhydrous THF were added *tert*-butanol (1.33 mL, 14.0 mmol). The resulting mixture was treated with DCC (412 mg, 2.0 mmol), followed by DMAP (240 mg, 2.0 mmol) at ambient temperature. Reaction mixture was stirred for 24 hours, then solids were removed by filtration. The filtrate was concentrated under reduced pressure and purified on silica gel (0-20% EtOAc-hexanes gradient) to give 385 mg (1.61 mmol, 81% yiled) of the title compound **17** as a colorless solid. $^1$H NMR (600 MHz, $CDCl_3$) δ 11.24 (s, 1H), 7.93 (d, *J* = 8.7 Hz, 1H), 7.76 (d, *J* = 2.3 Hz, 1H), 7.65 (dd, *J* = 8.7, 2.3 Hz, 1H), 1.64 (s, 9H). $^{13}$C NMR (151 MHz, $CDCl_3$) δ 168.51, 162.25, 151.91, 131.48, 118.80, 113.29, 113.02, 84.86, 28.22 (3C).

*(2S,3R,4S,5S,6S)-2-(2-(tert-butoxycarbonyl)-5-nitrophenoxy)-6-(methoxycarbonyl)tetrahydro-2H-pyran-3,4,5-triyl triacetate (**18**).* To a stirred solution of t-butyl 2-hydroxy-4-nitrobenzoate **17** (160 mg, 0.67 mmol) in 10 mL of anhydrous acetonitrile were added acetobromo-α-D-glucuronic acid methyl ester **6** (797 mg, 2.0 mmol). The resulting mixture was treated with silver (I) oxide (467 mg, 2.0 mmol) at room temperature in one portion, and stirred vigorously in the dark for 24 hours. Reaction mixture was diluted with 20 mL of EtOAc, filtered through a pad of silica gel, and concentrated under vacuum. The residue was purified on silica gel (EtOAc-hexanes, 0-30% v/v gradient) to obtain 220 mg (0.40 mmol, 61% yield) of compound **18** as a white solid. $^1$H NMR (600 MHz, CDCl3) δ 8.03 (d, *J* = 2.1 Hz, 1H), 7.94 (dd, *J* = 8.5, 2.1 Hz, 1H), 7.72 (d, *J* = 8.5 Hz, 1H), 5.43 – 5.38 (m, 1H), 5.37 – 5.29 (m, 3H), 4.29 (d, *J* = 9.3 Hz, 1H), 3.73 (s, 3H), 2.09 (s, 3H), 2.05 (s, 3H), 2.04 (s, 3H), 1.56 (s, 9H). $^{13}$C NMR (151 MHz, $CDCl_3$) δ 170.13, 169.43, 169.34, 166.72, 163.46, 155.01, 149.84, 131.08, 130.65, 118.02, 112.45, 99.39, 83.33, 72.69, 71.87, 70.93, 68.77, 53.17, 28.18 (3C), 20.78, 20.69, 20.64. LRMS (ESI): *m/z* 578.1 $[M+Na]^+$, Calcd for $C_{24}H_{29}NO_{14}$ *m/z* 578.2.

*(2S,3R,4S,5S,6S)-2-(2-(tert-butoxycarbonyl)-5-ureidophenoxy)-6-(methoxycarbonyl)tetrahydro-2H-pyran-3,4,5-triyl triacetate (**19**).* To a solution of nitro compound **18** (495 mg, 0.89 mmol) in MeOH (5 mL) were added Pd/C (10 wt. %, 200 mg). The flask was then evacuated and filled with $H_2$ gas from a balloon, in two repeating cycles. Reaction mixture was vigorously stirred for 3 days at room temperature with $H_2$ balloon attached. After the catalyst was removed by filtration through

a pad of celite, the filtrate was concentrated under vacuum to yield **19** (420 mg, 0.80 mmol, 90%) as an oil. The crude product was directly used for the next step without further purification. LRMS (ESI): *m/z* 526.7 $[M+H]^+$, Calcd for $C_{24}H_{31}NO_{12}$ *m/z* 526.2.

*Fmoc-AzaHIPS-4PN(Fmoc)-Sal-(o-Gluc-OAc)-OtBu (**20**).* To a solution of compound **19** (35 mg, 67 µmol) in DMF (1 mL) were added HIPS linker **15** (57 mg, 60 µmol), followed by DIPEA (32 µL, 0.18 mmol), and HATU (23 mg, 61 µmol). The resulting mixture was stirred for 1 hour at room temperature and then purified directly by reversed-phase chromatography (C18 column, 5-85% v/v acetonitrile-water with 0.05% TFA). Fractions containing the desired compound were pooled and concentrated under vacuum to give product **20** (49 mg, 34 µmol, 57 % yield) as a white solid. LRMS (ESI): *m/z* 1457.3 $[M+H]^+$, Calcd for $C_{79}H_{89}N_7O_{20}$ *m/z* 1456.6.

*Fmoc-AzaHIPS-4PN(Fmoc)-PEG2-Sal-(o-Gluc-OAc)-OH (**21**).* To a solution of *tert*-butyl ester **20** (49 mg, 34 µmol) in anhydrous DCM (0.5 mL) were added $SnCl_4$ in DCM (1 M, 0.34 mL, 0.34 mmol) at 0 °C. After stirring for 30 min, reaction mixture was quenched by addition of $H_2O$ (0.1 mL) and $CH_3CN$ (0.2 mL). The mixture was then concentrated under vacuum, and the residue was purified by reversed-phase chromatography (C18, 5-75% v/v $CH_3CN$-$H_2O$ with 0.05% TFA). Fractions containing the desired compound were pooled and concentrated under vacuum to give carboxylic acid **21** (40 mg, 29 µmol, 85% yield) as an oil. LRMS (ESI): *m/z* 1400.4 $[M+H]^+$, Calcd for $C_{75}H_{81}N_7O_{20}$ *m/z* 1400.6.

*H-Val-Cit-PABC-MMAE (**23**).* In an oven-dried 20 mL glass scintillation vial were mixed MMAE as a TFA salt (**13**, 125 mg, 0.15 mmol) and DIPEA (52 uL, 0.30 mmol) in 2 mL of anhydrous DMF. The resulting mixture was treated with PNP carbonate **22** (115 mg, 0.15 mmol) as a solid in portion at room temperature. Reaction mixture was stirred for 2 hours, then treated with 0.30 mL (3 mmol) of piperidine. After 30 minutes, reaction mixture was purified directly by reversed-phase chromatography (C18, acetonitrile-water 0-50% v/v gradient with 0.05% TFA). Pure fractions were evaporated under vacuum to give 123 mg (0.11 mmol, 73% yield over 2 steps) of the title compound **23** as a colorless solid. LRMS (ESI): *m/z* 1123.7 $[M+H]^+$, Calcd for $C_{58}H_{94}N_{10}O_{12}$ *m/z* 1123.7.

*AzaHIPS-4PNH-PEG2-Sal-(o-Gluc)-Val-Cit-PABC-MMAE (**2**).* To a solution of carboxylic acid **21** (15 mg, 10.7 µmol) in DMF (0.75 mL) were added H-Val-Cit-PABC-MMAE **23** (12 mg, 10.7 µmol) in DMA (0.5 mL), and DIPEA (6 µL, 34 µmol), followed by HATU (5 mg, 13 µmol). Reaction mixture was stirred for 1 hour at room temperature, then purified by reversed-phase HPLC (C18, acetonitrile-water 5-90% v/v gradient with 0.05% TFA). Fractions containing the desired compound were pooled and lyophilized to obtain protected precursor to compound **2** (5.2 mg, 2.1 µmol, 20% yield) as a white solid. LRMS (ESI): *m/z* 1253.6 $[M+2H]^{++}$, Calcd for $C_{133}H_{173}N_{17}O_{31}$ *m/z* 1253.1.

To a solution of protected precursor to construct **2** (5.2 mg, 2.1 µmol) in THF (0.2 mL) at 0 °C was slowly added a solution of lithium hydroxide (1 mg, 24 µmol) in 0.1 mL of $H_2O$. The resulting mixture was stirred at room temperature for 2 hours, then purified by reversed-phase HPLC (C18, acetonitrile-water 5-75% v/v gradient with 0.05% TFA). Fractions containing the desired compound were pooled and lyophilized to yield P3-glucuronide-MMAE construct **2** (2.5 mg, 1.3 µmol, 62% yield) as a white solid. HRMS (ESI): *m/z* 1921.0636 $[M+H]^{+}$, Calcd for $C_{96}H_{145}N_{17}O_{24}$ *m/z* 1921.0721.

##### Synthesis of control MMAE construct 3

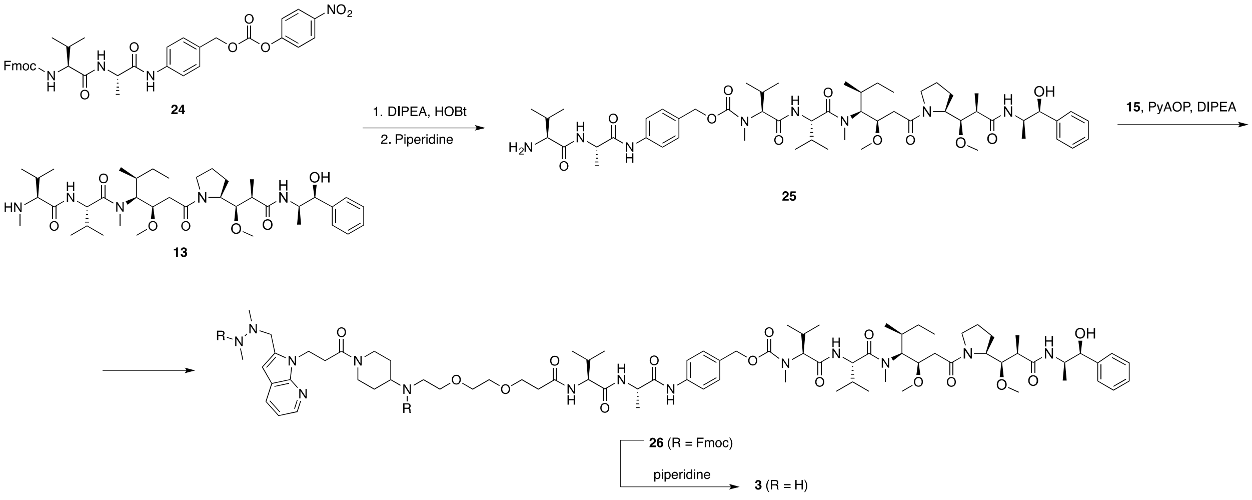


*H-Val-Ala-PABC-MMAE (**25**).* In an oven-dried 20 mL glass scintillation vial were mixed MMAE (**13**, 50 mg, 70 µmol), DIPEA (24 µL, 140 µmol), and HOBt (7 mg, 70 µmol) in 2 mL of anhydrous DMF. To this mixture were added PNP carbonate **24** (48 mg, 70 µmol) in one portion at room temperature. Reaction mixture was stirred for one hour, then piperidine (100 µL, 1 mmol) was added, and stirring was continued for 20 minutes. Reaction mixture was purified by reversed-phase chromatography (C18, 0-70% v/v $CH_3CN$-$H_2O$ with 0.05% TFA), fractions containing product were combined, concentrated under vacuum, and lyophilized to give product μ (45 mg, 43 μmol, 61% yield) as a white powder. LRMS (ESI): *m/z* 1037.7 $[M+H]^+$, Calcd for $C_{55}H_{88}N_8O_{11}$ *m/z* 1037.7.

*Fmoc-AzaHIPS-4PN(Fmoc)-PEG2-Val-Ala-PABC-MMAE (**26**).* To a stirred solution of amine **25** (45 mg, 43 μmol) and HIPS linker **15** (41 mg, 43 μmol) in 2 mL of DMF were added DIPEA (23 μL, 0.13 mmol), followed by PyAOP (23 mg, 43 μmol) in one portion at room temperature. The resulting mixture was stirred for 30 minutes, then purified directly by reversed-phase chromatography (C18, 0-100% v/v $CH_3CN$-$H_2O$ with 0.05% TFA). Pure fractions were pooled, concentrated, and lyophilized to afford compound **26** (71 mg, 36 μmol, 84% yield) as a white solid. LRMS (ESI): *m/z* 985.2 $[M+2H]^{++}$, Calcd for $C_{110}H_{146}N_{14}O_{19}$ *m/z* 984.6.

*AzaHIPS-4PNH-PEG2-Val-Ala-PABC-MMAE (**3**).* To a stirred solution of compound **26** (71 mg, 36 μmol) in 3 mL of DMA were added 70 μL of piperidine (0.72 mmol) at room temperature. Reaction mixture was stirred for 30 minutes until Fmoc deprotection was judged complete by LCMS analysis, then directly purified by reversed-phase HPLC (C18, 0-50% v/v $CH_3CN$-$H_2O$ with 0.05% TFA). Fractions were pooled and lyophilized to afford 43 mg of compound **3** (28 μmol, 78% yield) as a white powder. HRMS (ESI): *m/z* 1523.9555 $[M+H]^+$, Calcd for $C_{80}H_{126}N_{14}O_{15}$ *m/z* 1523.9600.

**Synthesis of control MMAE construct 4**

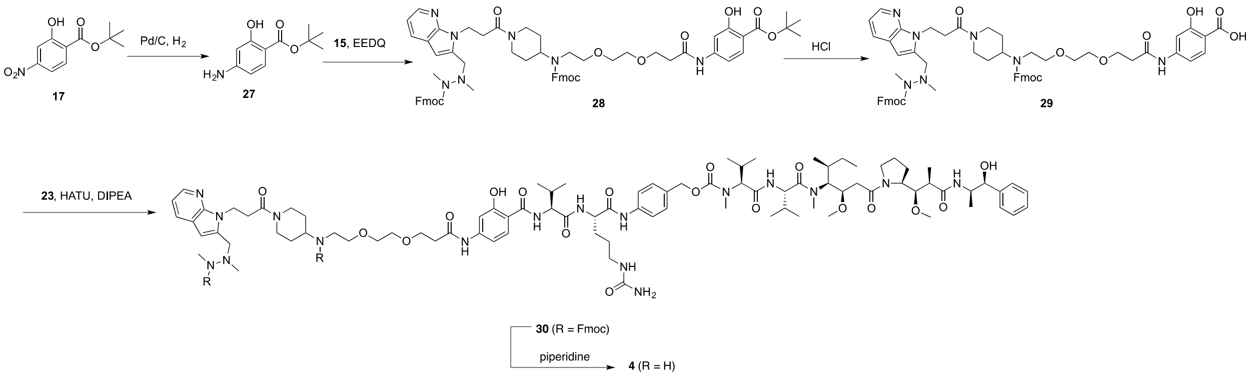

<u>*tert-Butyl 4-amino-2-hydroxybenzoate (27).*</u> To a solution of nitro compound **17** (485 mg, 2.0 mmol) in EtOAc (4 mL) and EtOH (4 mL) were added palladium on carbon (10 wt.%, 215 mg, 0.2 mmol). The flask was then evacuated and filled with $H_2$ gas from a balloon, in two repeating cycles. Reaction mixture was vigorously stirred for 2 days at room temperature with $H_2$ balloon attached. After the catalyst was removed by filtration through a pad of celite, the filtrate was concentrated under vacuum to yield **27** (418 mg, 2 mmol, quant. yield) as an oil, which was directly used in the next step without further purification. LRMS (ESI): *m/z* 210.1 $[M+H]^+$, Calcd for $C_{11}H_{15}NO_3$ *m/z* 210.1.

<u>*Fmoc-AzaHIPS-4PN(Fmoc)-PEG2-Sal-COOtBu (28).*</u> To a mixture of aniline **27** (16 mg, 77 µmol) and HIPS linker **15** (47 mg, 50 µmol) in THF (0.5 mL) were added EEDQ (25 mg, 0.10 mmol). The resulting mixture was stirred for one hour at room temperature, the purified by reversed-phase chromatography (C18, 5-85% v/v $CH_3CN$-$H_2O$ with 0.05% TFA). Fractions containing the desired compound were pooled and concentrated under vacuum to yield compound **28** (18 mg, 16 µmol, 32% yield) as a white solid. LRMS (ESI): *m/z* 1140.4 $[M+H]^+$, Calcd for $C_{66}H_{73}N_7O_{11}$ *m/z* 1140.5.

<u>*Fmoc-AzaHIPS-4PN(Fmoc)-PEG2-Sal-COOH (29).*</u> To a solution of tert-butyl ester **28** (18mg, 16 µmol) in dioxane (0.1 mL) at 0 °C were added HCl in dioxane (4 M, 0.1 mL, 0.4 mmol). The reaction mixture was warmed up to room temperature and stirred for one hour, then purified by reversed-phase chromatography (C18, 5-85% v/v $CH_3CN$-$H_2O$ with 0.05% TFA) to yield carboxylic acid **29** (9 mg, 8 µmol, 50 % yield) as a yellow oil. LRMS (ESI): *m/z* 1084.4 $[M+H]^+$, Calcd for $C_{62}H_{65}N_7O_{11}$ *m/z* 1084.5.

<u>*Fmoc-AzaHIPS-4PN(Fmoc)-PEG2-Sal-Val-Cit-PABC-MMAE (30).*</u> To a solution of carboxylic acid **29** (12 mg, 11 µmol) in DMF (50 µL) were added H-Val-Cit-PABC-MMAE **23** (12 mg, 11 µmol), DIPEA (6 µL, 34 µmol), followed by HATU (5 mg, 13 µmol). Reaction mixture was stirred for one hour at room temperature, then purified by reversed-phase HPLC (C18, 5-90% v/v $CH_3CN$-$H_2O$ with 0.05% TFA). Fractions containing the desired product were pooled and lyophilized to yield compound **30** (3 mg, 13 % yield) as a white solid. LRMS (ESI): *m/z* 1095.6 $[M+2H]^{++}$, Calcd for $C_{120}H_{157}N_{17}O_{22}$ *m/z* 1095.1.

*AzaHIPS-4PNH-PEG2-Sal-Val-Cit-PABC-MMAE (**4**).* To a solution of **30** (2.8 mg, 1.3 µmol) in DMA (0.2 mL) at 0 °C was slowly added piperidine (1.0 µL, 10 µmol). The resulting mixture was stirred for 30 min at room temperature. then purified by reversed-phase HPLC (C18, 5-70% v/v $CH_3CN$-$H_2O$ with 0.05% TFA).The fractions containing the desired compound were pooled and lyophilized to yield **4** (1.0 mg, 45% yield) as a white solid. HRMS (ESI): *m/z* 1745.0332 $[M+H]^+$, Calcd for $C_{90}H_{137}N_{17}O_{18}$ *m/z* 1645.0400.

**Synthesis of HIPS linker 15**

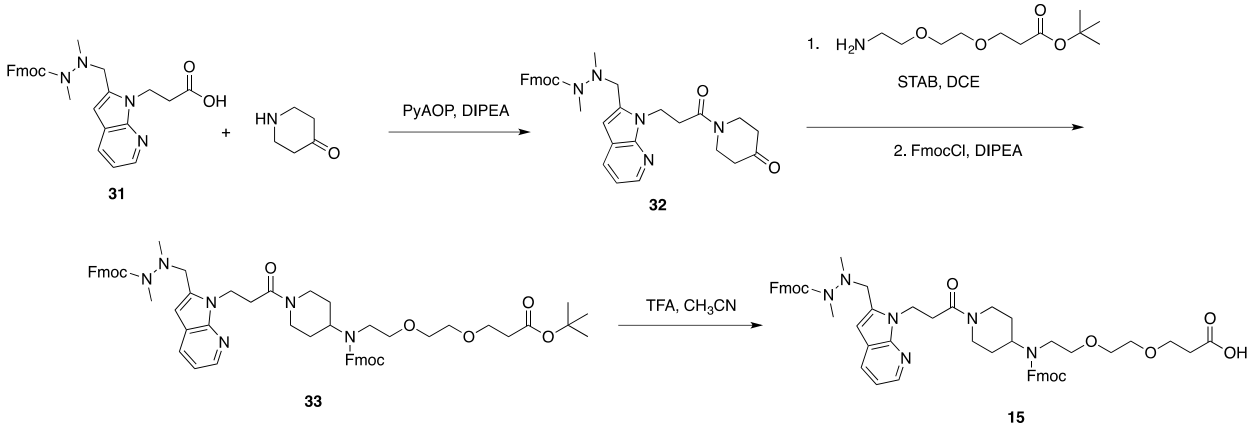


*(9H-fluoren-9-yl)methyl 1,2-dimethyl-2-((1-(3-oxo-3-(4-oxopiperidin-1-yl)propyl)-1H-pyrrolo[2,3-b]pyridin-2-yl)methyl)hydrazine-1-carboxylate (**32**).* In a round-bottom flask were mixed Fmoc-AzaHIPS-COOH **31** (2.5 g, 5.2 mmol), 4-piperidinone (hydrochloride monohydrate, 0.85 g, 8.6 mmol), and DIPEA (2.5 mL, 14.4 mmol) in 12 mL of anhydrous DMF. The mixture was stirred and treated with PyAOP (2.9 g, 5.6 mmol) at room temperature in small portions. Reaction mixture was stirred for 45 minutes and poured into 100 mL of 10% aqueous citric acid solution, and extracted with ethyl acetate (2x75 mL). Extracts were washed with brine and dried over sodium sulfate. After removal of solvent under vacuum, the residue was purified on silica gel (0-5% MeOH-EtOAc) to afford 2.5 g (4.4 mmol, 82 % yield) of compound **32** as a yellowish solid foam. LRMS (ESI): *m/z* 565.9 $[M+H]^+$, Calcd for $C_{33}H_{35}N_5O_4$ *m/z* 566.3.

*Fmoc-AzaHIPS-4PN(Fmoc)-PEG2-OtBu (**33**).* To an oven-dried 50 mL round-bottom flask were loaded ketone 15b (2.5g, 4.4 mmol) and $NH_2$-PEG2-OtBu (1.1 g, 4.7 mmol), followed by 15 mL of anhydrous 1,2-dichloroethane under nitrogen. The resulting mixture was stirred for 30 minutes at room temperature, then treated with sodium triacetoxyborohydride (1.9 g, 9 mmol) in one portion at room temperature. Reaction mixture was stirred overnight, diluted with 100 mL of DCM and washed once with saturated aqueous sodium bicarbonate solution (50 mL). Organic layer was dried over sodium sulfate, concentrated under vacuum, and the residue was dissolved in 15 mL of anhydrous DCM. To this solution were added DIPEA (2.3 mL, 13 mmol) and Fmoc chloride (1.7 g, 6.6 mmol). Reaction mixture was stirred for one hour until the reaction was judged complete by LCMS analysis, diluted with 100 mL of DCM, washed sequentially with 10% aqueous citric acid (50 mL), water, and brine, and dried over sodium sulfate. After removal of solvent, the residue was purified on silica gel (1-4% v/v MeOH-DCM) to give 2.4 g of product **33** (2.4 mmol, 55% yield over two steps). LRMS (ESI): *m/z* 1004.9 $[M+H]^+$, Calcd for $C_{59}H_{68}N_6O_9$ *m/z* 1005.1.

*Fmoc-AzaHIPS-4PN(Fmoc)-PEG2-COOH (**15**).* *tert*-Butyl ester **33** (2.4 g, 2.4 mmol) was dissolved in 4 mL of anhydrous acetonitrile. To this solution were added 16 mL of TFA, and the resulting mixture was stirred at room temperature for 30 minutes, concentrated under reduced pressure, and purified directly by reversed-phase flash chromatography (C18, 0-100% v/v $CH_3CN$-$H_2O$). Fractions containing the desired product were combined, concentrated under vacuum, and partitioned between brine (50 mL) and ethyl acetate (75 mL). Aqueous layer was extracted with ethyl acetate (2x25 mL), combined organic layers were dried over sodium sulfate. Removal of solvent afforded 2.0 g (2.1 mmol, 88% yield) of the title compound **15** as an off-white solid. $^1H$ NMR (600 MHz, DMSO) δ 8.25 (s, 1H), 7.99 – 7.81 (m, 4H), 7.80 – 7.66 (m, 2H), 7.65 – 7.48 (m, 4H), 7.43 – 7.20 (m, 8H), 7.12 (dd, *J* = 7.8, 4.9 Hz, 1H), 4.73 – 4.11 (m, 10H), 3.80 – 3.43 (m, 5H), 3.43 – 3.33 (m, 3H), 3.31 – 3.12 (m, 2H), 3.08 – 2.75 (m, 6H), 2.68 (s, 3H), 2.65 – 2.54 (m, 1H), 2.46 – 2.28 (m, 3H), 2.18 – 2.02 (m, 2H), 1.55 – 1.26 (m, 2H), 1.08 – 0.73 (m, 2H). $^{13}C$ NMR (151 MHz, DMSO) δ 172.58, 172.00, 170.33, 168.39, 158.73, 158.48, 158.23, 157.98, 154.92, 144.05, 140.94, 127.59, 127.50, 127.40, 127.09, 127.02, 124.83, 124.64, 120.10, 118.18, 116.26, 115.72, 114.34, 112.42, 100.79, 69.58, 69.50, 68.91, 68.63, 66.31, 66.24, 65.83, 59.75, 54.48, 46.90, 46.66, 44.25, 40.49, 39.94, 39.80, 39.66, 39.52, 39.38, 39.24, 39.10, 38.61, 34.72, 32.34, 29.34, 28.92, 28.53, 21.04, 20.74, 14.07. LRMS (ESI): *m/z* 948.8 $[M+H]^+$, Calcd for $C_{55}H_{60}N_6O_9$ *m/z* 949.4.

### NMR Spectra

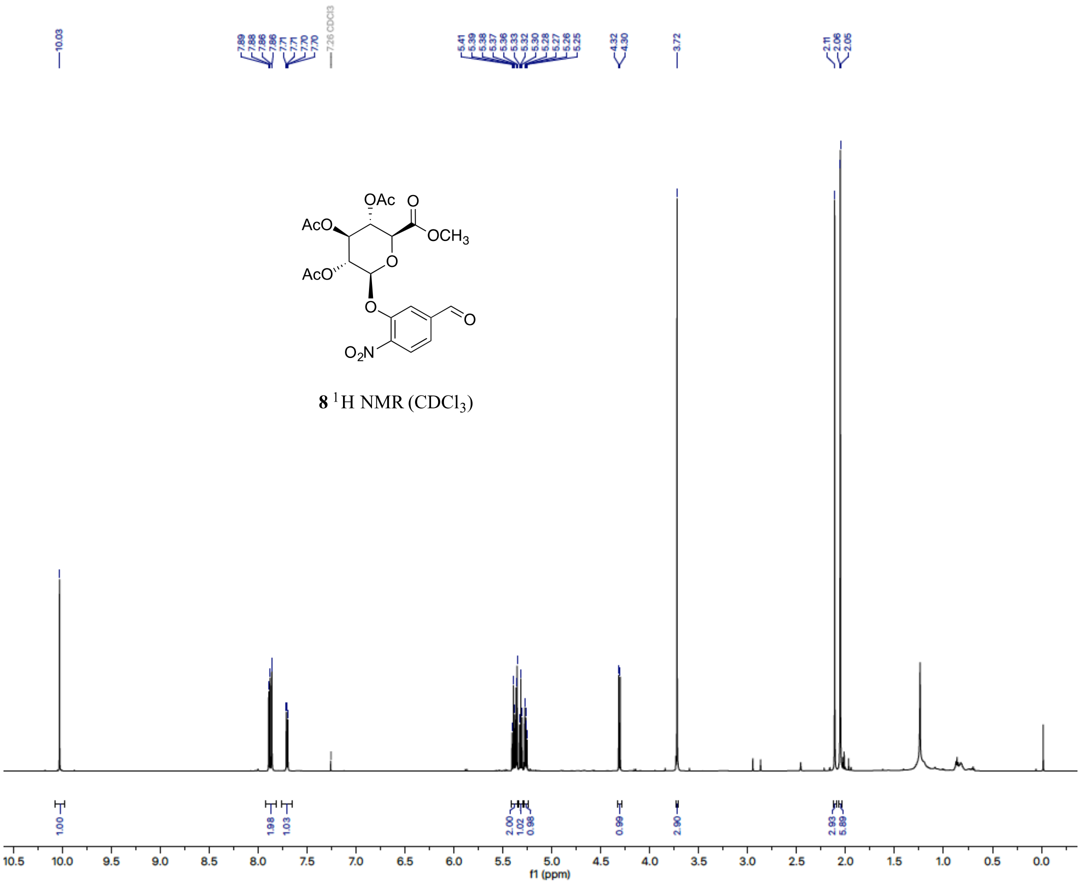

**8** [1]H NMR ($CDCl_3$)

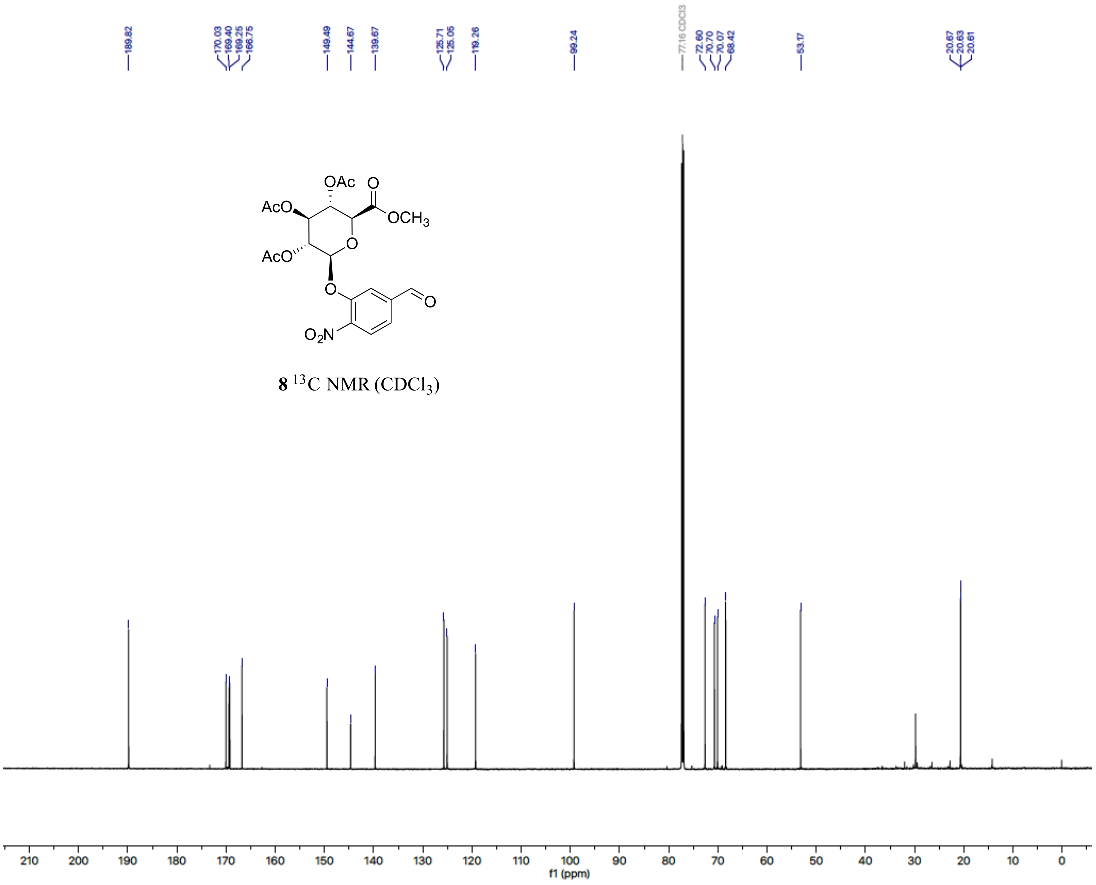

**8** [13]C NMR ($CDCl_3$)

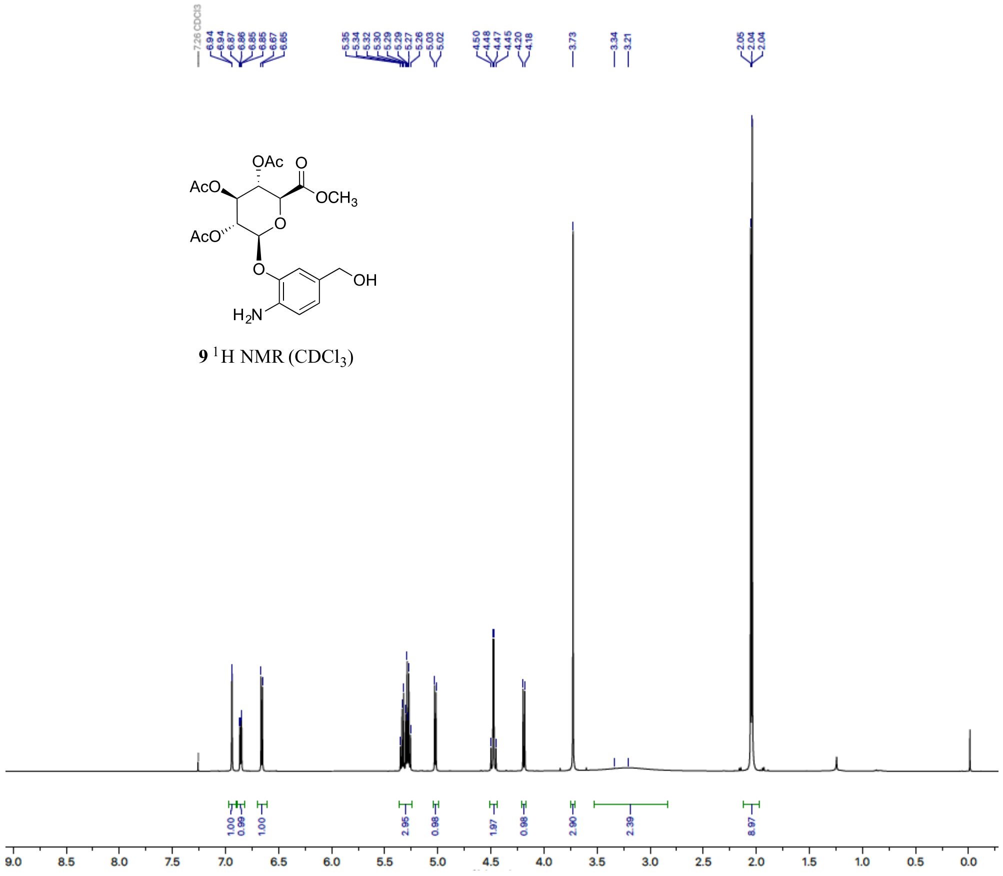
OAc
O
AcO
OCH3
AcO
O
O
OH
H2N
9 1H NMR (CDCl3)

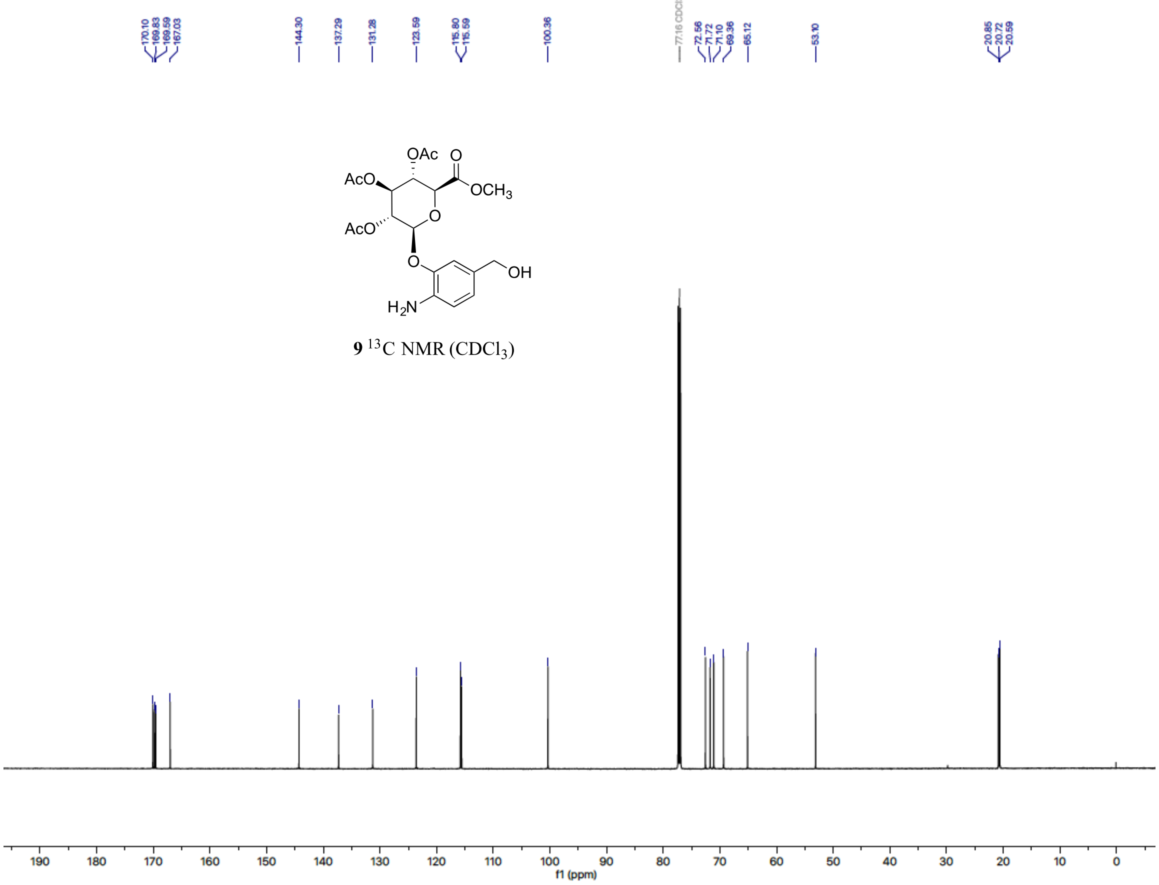
OAc
O
AcO
OCH3
AcO
O
O
OH
H2N
9 13C NMR (CDCl3)
f1 (ppm)

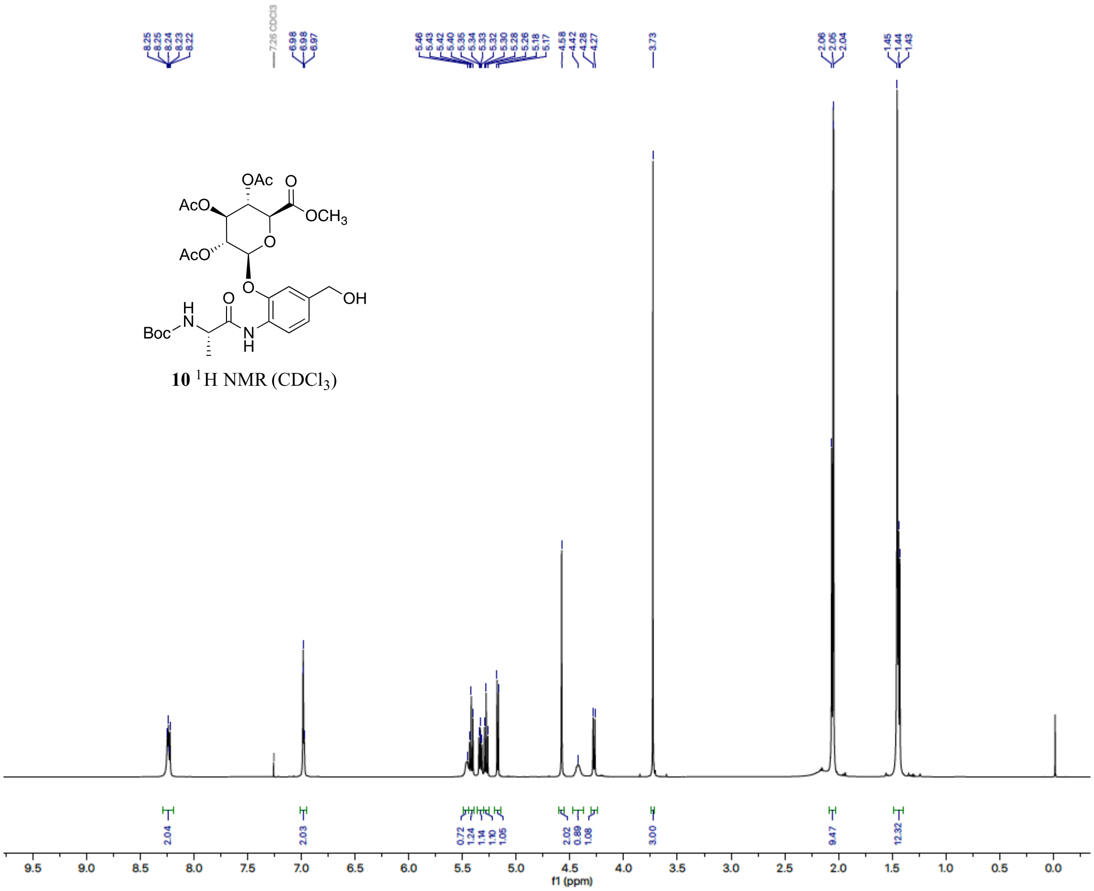

**10** $^{1}$H NMR ($CDCl_3$)

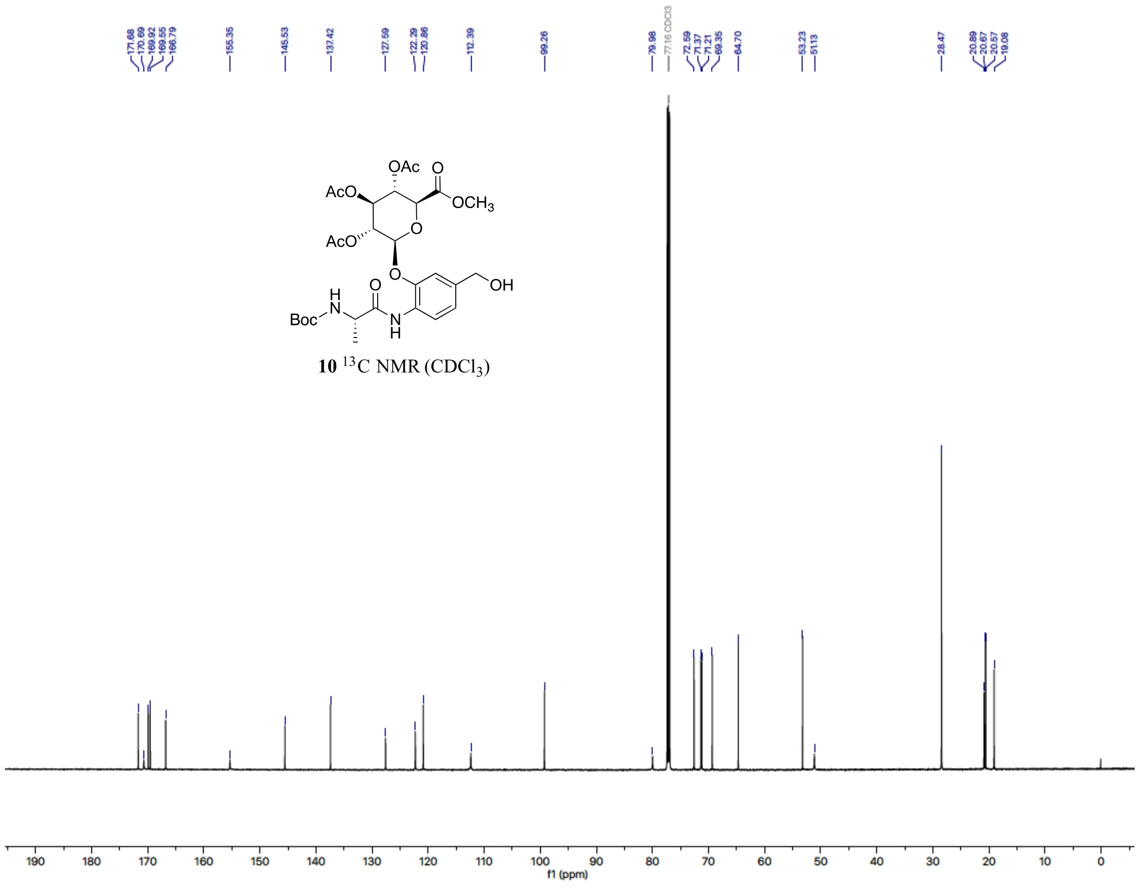

**10** $^{13}$C NMR ($CDCl_3$)

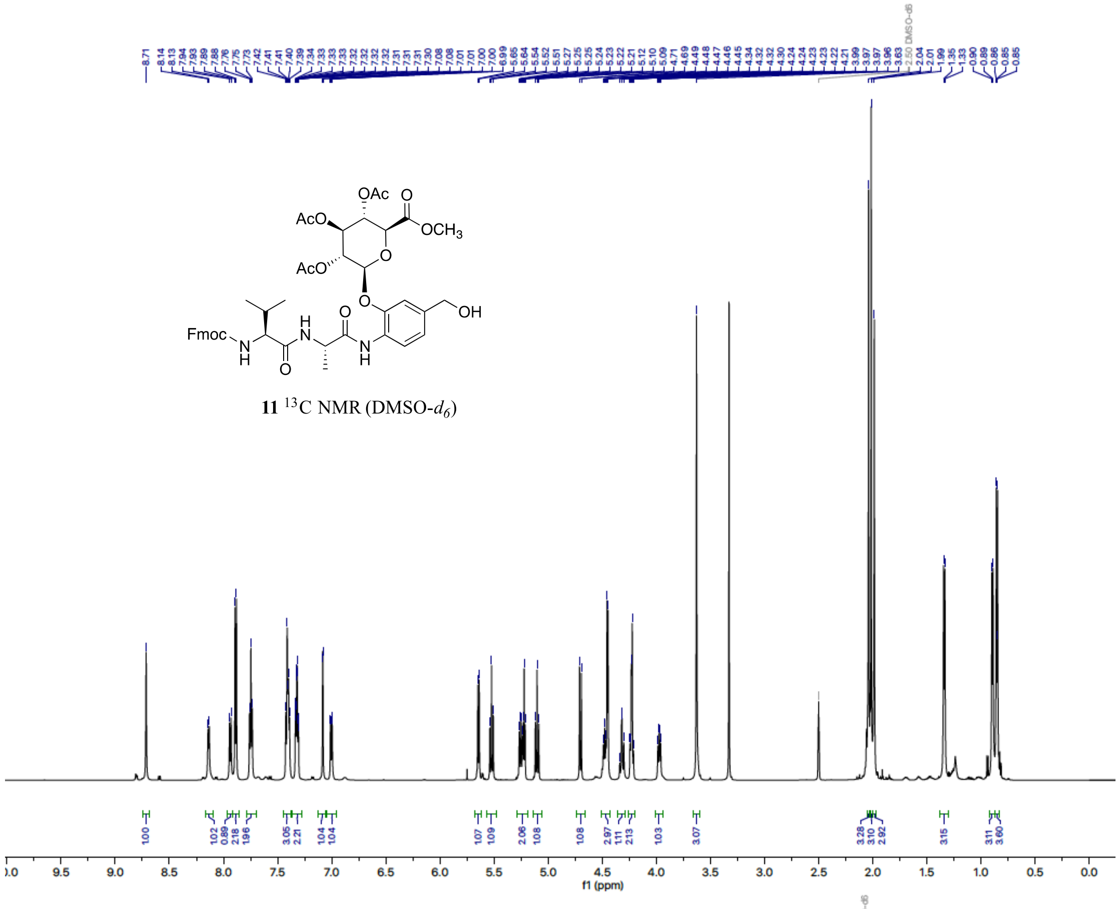


**11** $^{13}$C NMR (DMSO-$d_6$)

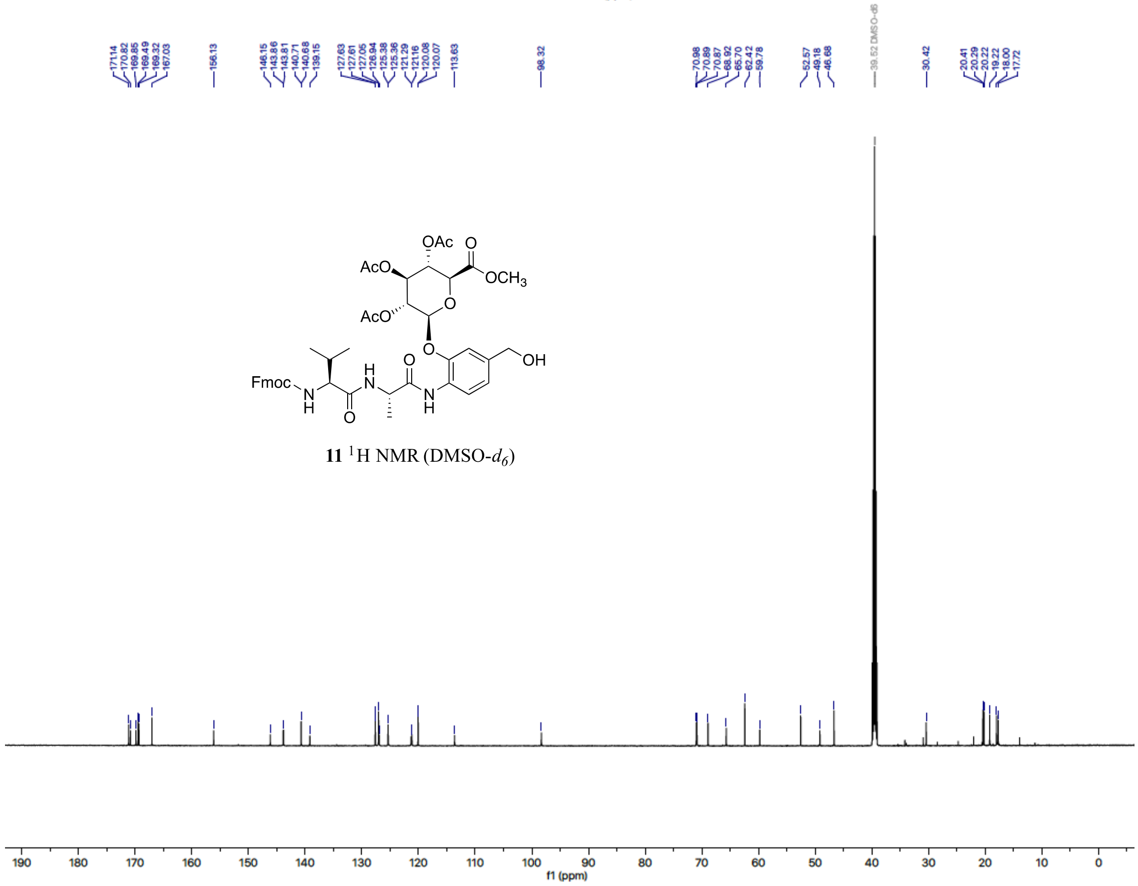


**11** $^{1}$H NMR (DMSO-$d_6$)

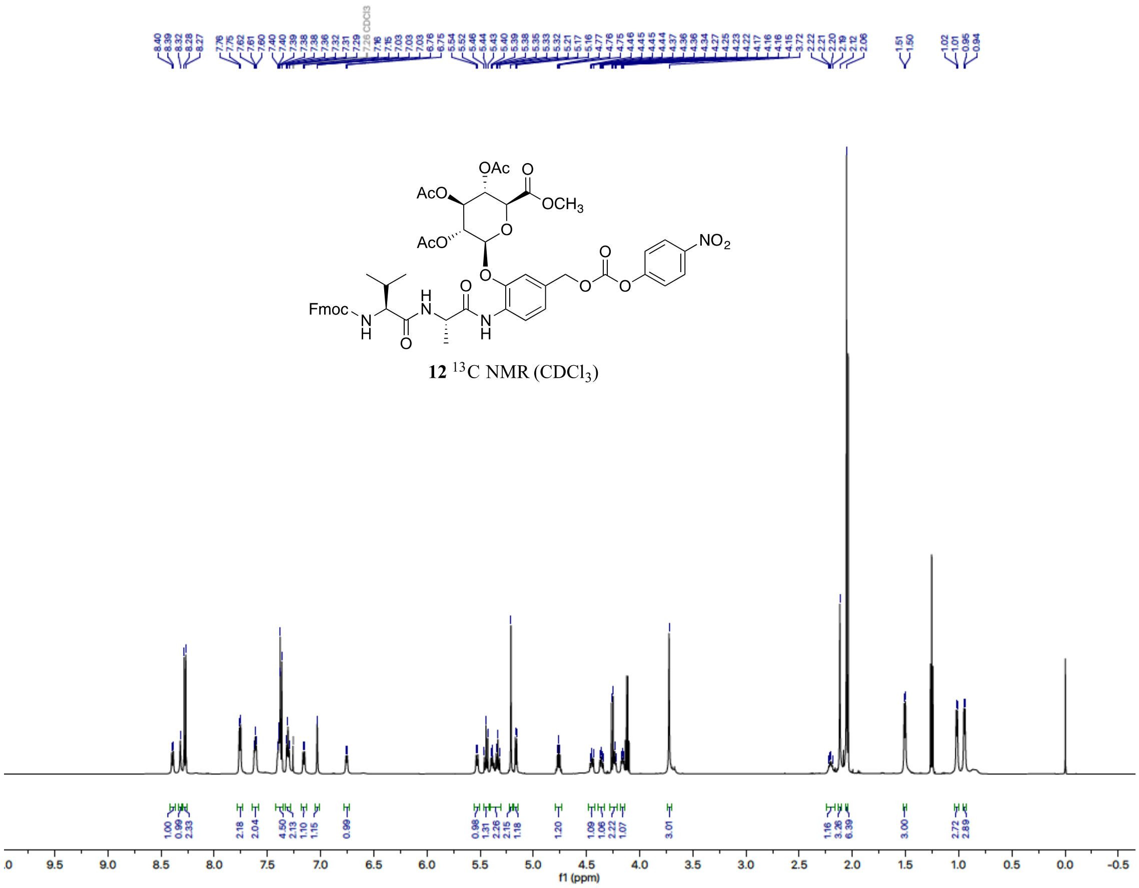


**12** $^{13}$C NMR ($CDCl_3$)

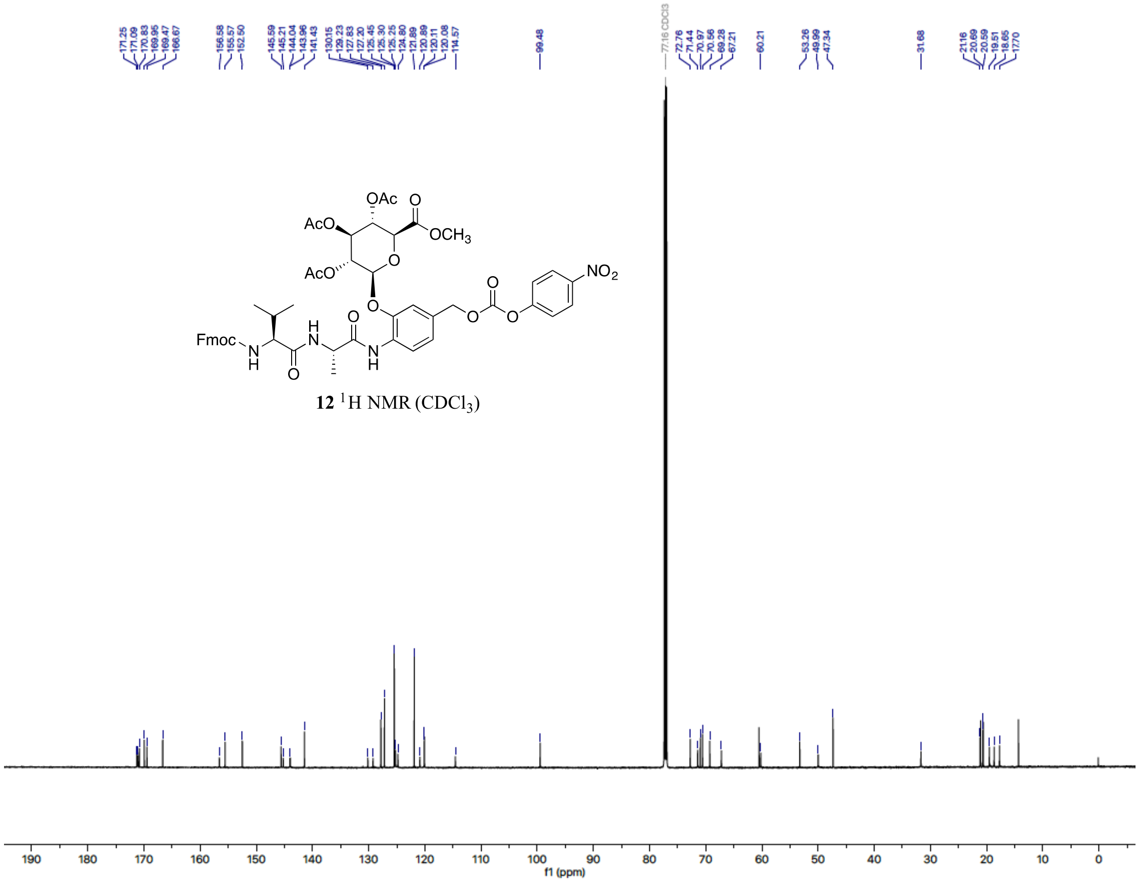


**12** $^{1}$H NMR ($CDCl_3$)

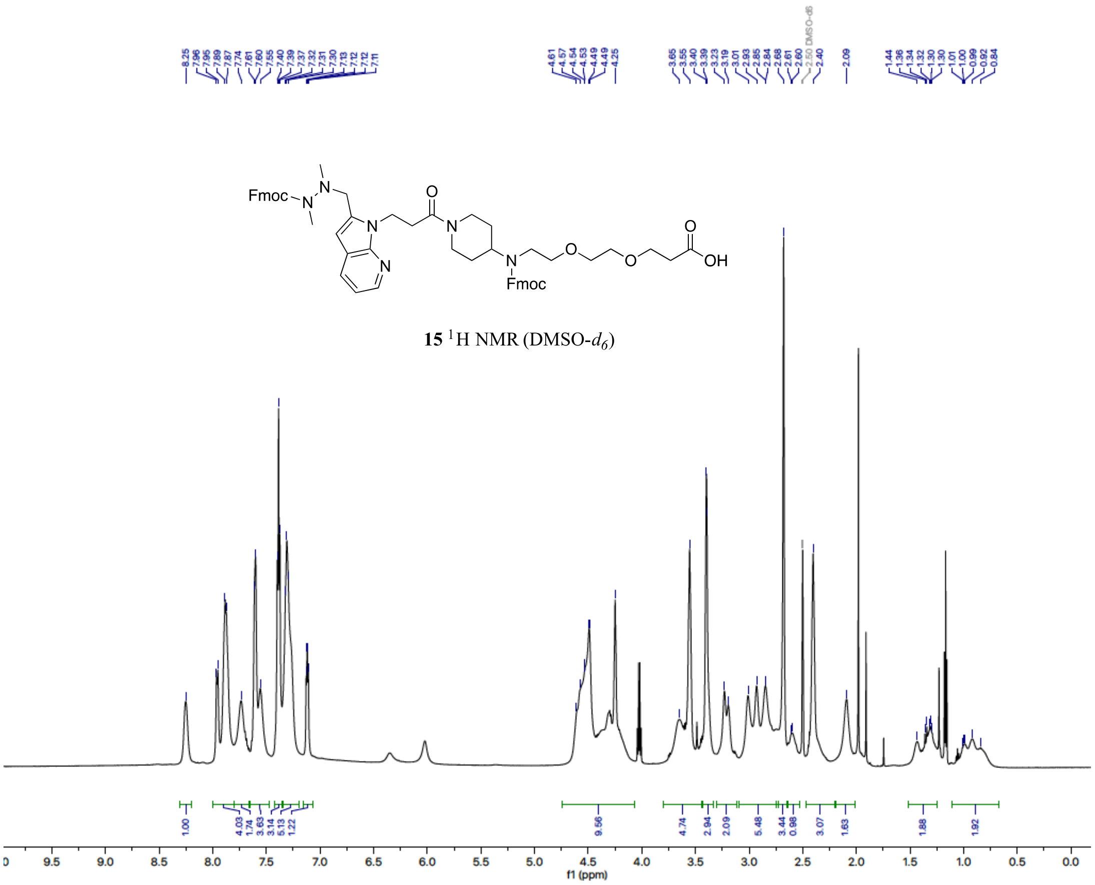


**15** $^{1}$H NMR (DMSO-$d_6$)

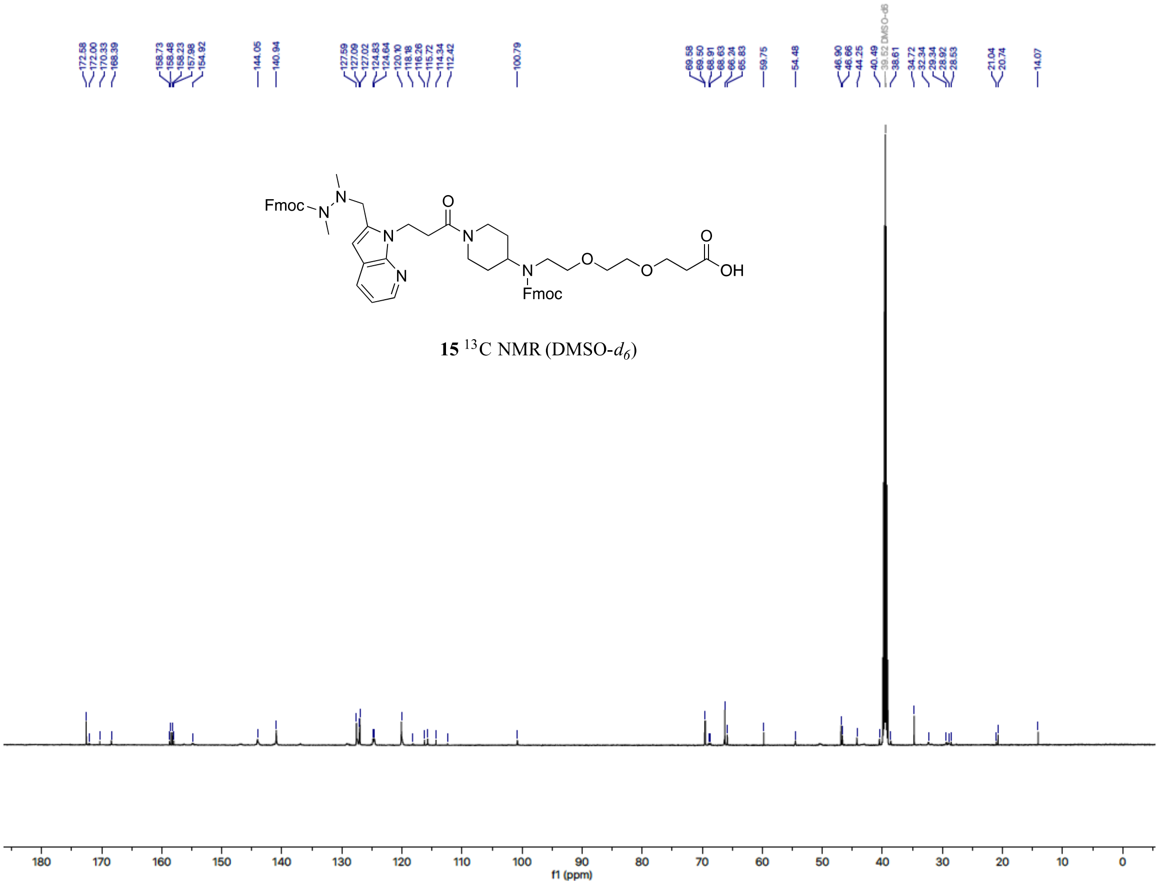


**15** $^{13}$C NMR (DMSO-$d_6$)

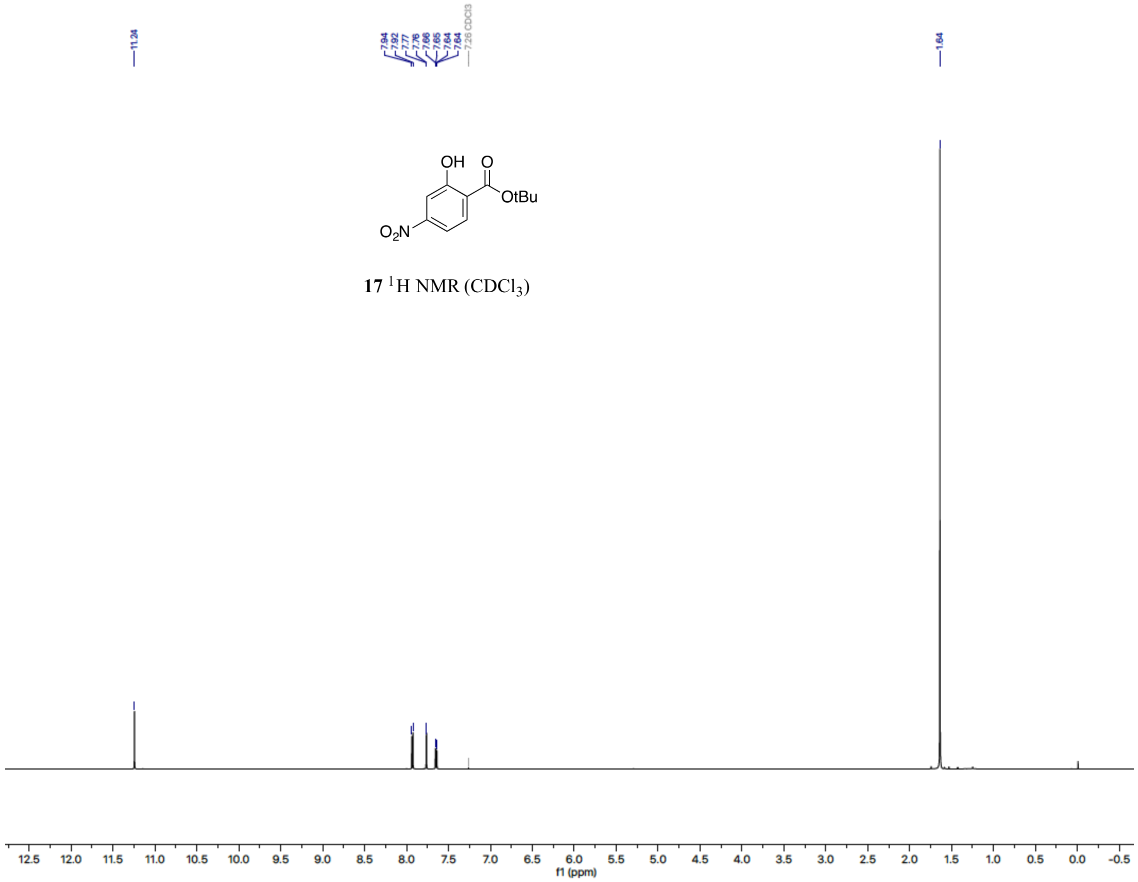
11.24
7.94
7.92
7.77
7.76
7.66
7.65
7.64
7.64
7.26 CDCl3
1.64
OH
O
OtBu
$O_2N$
**17** $^{1}$H NMR ($CDCl_3$)
f1 (ppm)

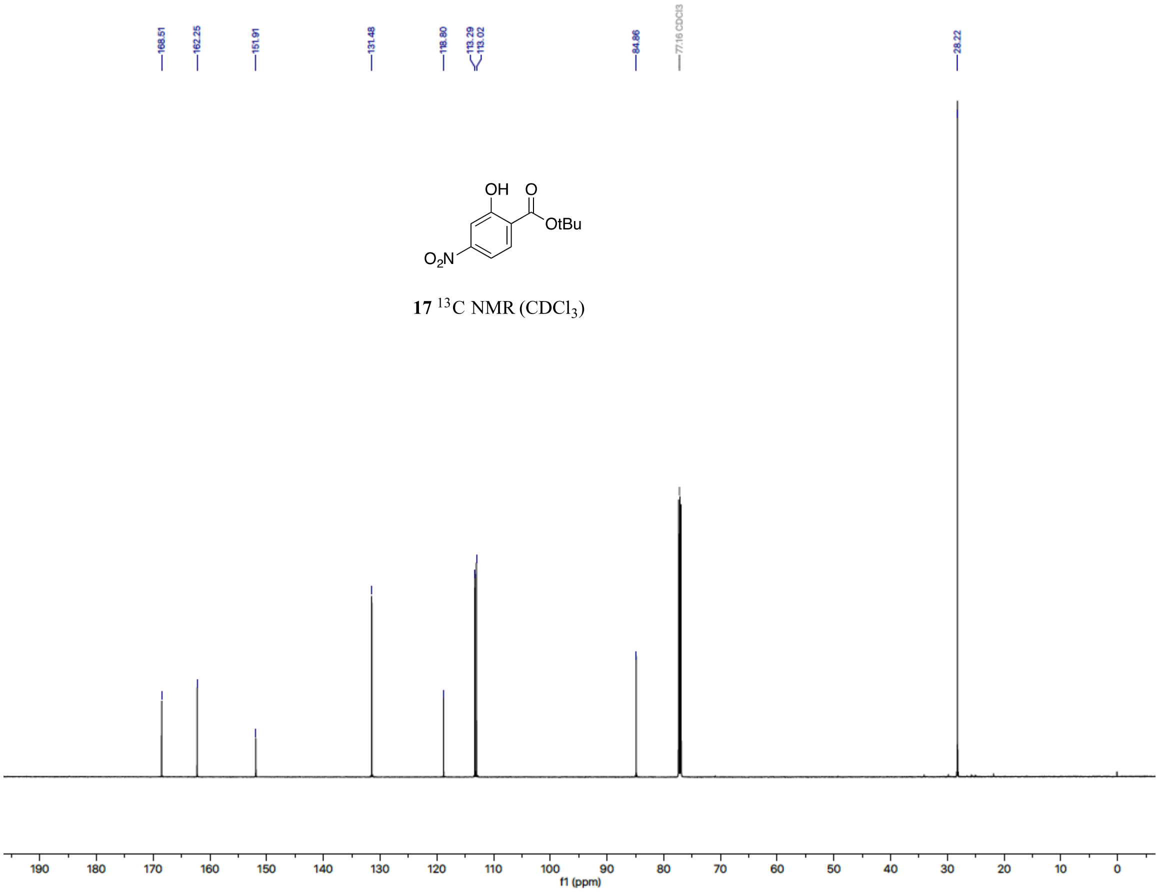
168.51
162.25
151.91
131.48
118.80
113.29
113.02
84.86
77.16 CDCl3
28.22
OH
O
OtBu
$O_2N$
**17** $^{13}$C NMR ($CDCl_3$)
f1 (ppm)

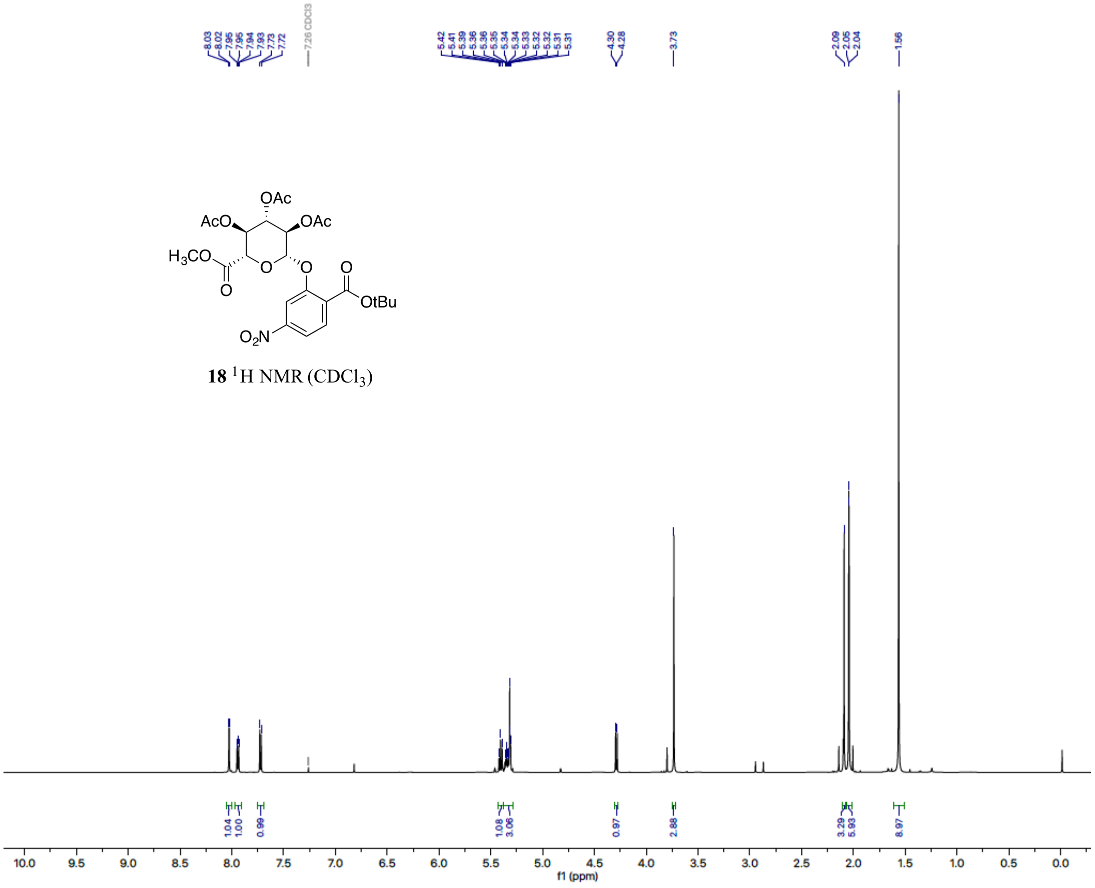


**18** $^{1}$H NMR ($CDCl_3$)

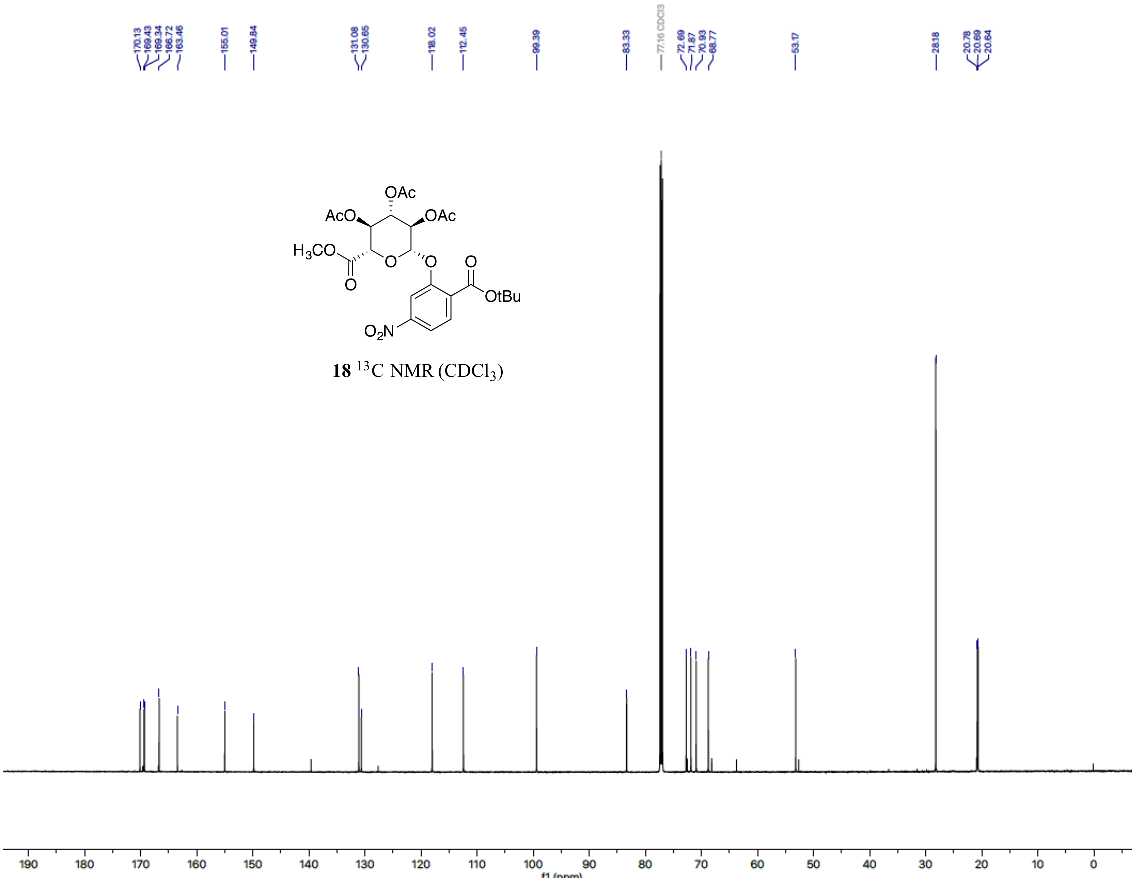


**18** $^{13}$C NMR ($CDCl_3$)

**Table SI-1**. DARs of ADCs used in the reported studies**.**

| ADC Composition | ADC Lot | DAR | Study type |
|---|---|---|---|
| 1-CD79b | cRW2725 | 1.74 | in vitro potency |
| 2-CD79b | cRW2727 | 1.66 | in vitro potency |
| 3-CD79b | cRW2595 | 1.82 | in vitro potency |
| 4-CD79b | cRW2726 | 1.4 | in vitro potency |
| 5-CD79b | cRW2714 | 3.18 | in vitro potency |
| 1-CD79b | cRW1468 | 1.73 | in vitro stability |
| 2-CD79b | cRW1471 | 1.67 | in vitro stability |
| 3-CD79b | cRW2595 | 1.82 | in vitro stability |
| 4-CD79b | cRW1470 | 1.52 | in vitro stability |
| 5-CD79b | cRW2587 | 2.62 | in vitro stability |
| 1-CD79b | cRW1504A | 1.75 | xenograft |
| 2-CD79b | cRW1510B | 1.8 | xenograft |
| 3-CD79b | cRW1505B | 1.67 | xenograft |
| 5-CD79b | cRW1326 | 3.5 | xenograft |
| 1-CD79b | cRW2286 | 1.75 | rat tox |
| 3-CD79b | cRW2285 | 1.71 | rat tox |
| 5-CD79b | cRW2312 | 3.47 | rat tox |
